# dsRNA-induced immunity targets plasmodesmata and is suppressed by viral movement proteins

**DOI:** 10.1101/2022.11.21.517408

**Authors:** Caiping Huang, Ana Rocio Sede, Laura Elvira-González, Yan Yan, Miguel Rodriguez, Jerome Mutterer, Emmanuel Boutant, Libo Shan, Manfred Heinlein

## Abstract

Emerging evidence indicates that in addition to the well-recognized antiviral RNA silencing, dsRNA elicits responses of pattern-triggered immunity (PTI), likely contributing plant resistance against virus infections. However, compared to bacterial and fungal elicitor-mediated PTI, the mode-of-action and signaling pathway of dsRNA-induced defense remain poorly characterized. Here, using multi-color *in vivo* imaging by GFP mobility, staining of callose and plasmodesmal marker lines, we show that dsRNA-induced PTI restricts the progression of virus infection by triggering callose deposition at plasmodesmata, thereby likely limiting the macromolecular transport through these cell-to-cell communication channels. The plasma membrane-resident kinase module of SERK1 and BIK1/PBL1, plasmodesmata-localized proteins PDLP1/2/3 and calmodulin-like CML41, and Ca^2+^ signals are involved in the dsRNA-induced signaling leading to callose deposition at plasmodesmata and antiviral defense. In addition, unlike classical bacterial elicitor flagellin, dsRNA does not trigger detectable reactive oxygen species (ROS) burst, further substantiating a partially shared immune signaling framework with distinct features triggered by different microbial patterns. Likely as a counteract strategy, viral movement proteins from different viruses suppress the dsRNA-induced host response leading to callose deposition to achieve infection. Thus, our data support the new model of how plant immune signaling constrains the virus movement by inducing callose deposition at plasmodesmata and how viruses counteract this layer of immunity.

**One-sentence summary:** dsRNA-induced antiviral PTI targets plasmodesmata for callose deposition and is suppressed by virus-encoded movement proteins.

**IN A NUTSHELL:** *Background:* Plants use different defense mechanisms pathogens. The major mechanism that plants use for defense against viruses is known as RNA silencing. This mechanism is triggered by the presence of viral double-stranded (ds)RNA and uses small RNAs to inhibit viral replication by targeting the viral genome for degradation. Recently, it was found that dsRNA elicits antiviral defense also through a protein-mediated mechanism known as pattern-triggered immunity (PTI). However, the underlying mechanism of antiviral PTI and how viruses overcome this plant defense mechanism to cause infection is unknown.

*Question:* In this study we asked how dsRNA-induced PTI acts to inhibit virus infection and whether we can identify components of the PTI signaling pathway. Moreover, we wanted to know how viruses overcome this plant host defense response in order to cause infection.

*Findings:* We demonstrate that dsRNA-induced PTI targets plasmodesmata (PD), the intercellular communication conduits in plant cell walls that viruses use to spread infection from cell to cell. By inducing the deposition of callose, dsRNA-induced PTI reduces PD permeability, thus restricting virus movement. We identified PTI signaling components required for dsRNA-induced PD callose deposition and delineate a PTI pathway showing important difference to PTI pathways triggered by microbial elicitors. Moreover, viral movement proteins (MPs) suppress the dsRNA-induced callose deposition response at PD. This leads to a new model of how plant immune signaling constrains virus movement and how viruses counteract this layer of immunity.

*Next steps:* This study calls upon the identification of the PTI dsRNA receptor and the mechanisms of PTI signaling (involving identified components such as SERK1, BIK1, calcium channels, CML41, PDLP1/2/3) and PTI suppression by MPs, and how dsRNA-induced PTI and RNA silencing are controlled during the spread of infection.

## Introduction

The virome of plants is dominated by RNA viruses (Dolja et al., 2020) and several of these cause devastating diseases in cultivated plants leading to global crop losses (Jones and Naidu, 2019; Jones, 2021). To infect plants, RNA viruses engage in complex interactions with compatible plant hosts. In cells at the spreading infection front, RNA viruses associate with cellular membranes and replicate their genome through double-stranded RNA (dsRNA) intermediates. Moreover, they use their movement proteins (MP) to interact with membrane-associated transport processes in order to achieve the movement of replicated genome copies through cell wall nanochannels called plasmodesmata (PD) in order to infect new cells (Heinlein, 2015). Importantly, sensing of viral dsRNA by the host triggers defense responses against infection, which viruses must be able to control in order to propagate.

The most important antiviral host response in plants is RNA silencing (Ding and Voinnet, 2007). It involves host DICER-LIKE enzymes that inhibit viral replication by cleaving the viral dsRNA replication intermediate into small interfering RNAs (siRNAs). These viral siRNAs can associate with ARGONAUTE proteins in RNA-induced silencing complexes (RISCs) to further guide the sequence-specific degradation and translational suppression of viral RNA. To control this antiviral response and to enhance their replication, viruses have evolved specific effector proteins that interfere with the RNA silencing pathway at distinct steps (Csorba et al., 2015). More recent research has shown that in addition to the antiviral RNA silencing response, RNA virus infection also activates pattern-triggered immunity (PTI) (Kørner et al., 2013), whereby dsRNA acts as an important elicitor (Niehl et al., 2016). Unlike RNA silencing, PTI is triggered by specific recognition of conserved microbe- or pathogen-associated molecular patterns (MAMPs or PAMPs) by pathogen-recognition receptors (PRRs) and the induction of defense signaling (DeFalco and Zipfel, 2021). Importantly, dsRNA-induced PTI is independent of dsRNA sequence. Thus, PTI is activated by viral dsRNA but also as well by non-viral dsRNA, for example GFP dsRNA or the synthetic dsRNA analog polyinosinic-polycytidilic acid [poly(I:C)] (Niehl et al., 2016), a well-known ligand of the dsRNA-perceiving TLR3-receptor in animals (Alexopoulou et al., 2001). Similar to virus replication, treatment of *Arabidopsis thaliana* plants with poly(I:C) elicits antiviral defense along with activating typical PTI responses, such as mitogen-activated protein kinase (MPK), ethylene production, seedling root growth inhibition, and marker gene expression (Kørner et al., 2013; Niehl et al., 2016). Poly(I:C)-triggered ethylene production and antiviral defense were shown to depend on the co-receptor kinase SOMATIC EMBRYOGENESIS RECEPTOR-LIKE KINASE 1 (SERK1) (Niehl et al., 2016) but neither other components of the signaling pathway nor the mechanism by which PTI restricts virus infection are known. Here, we demonstrate that unlike RNA silencing, which controls viral RNA accumulation, dsRNA-induced PTI acts on PD to restrict virus movement. New components of the PTI signaling mechanism to PD are identified and shown to be critical for limiting virus infection and symptom formation. Moreover, the observations indicate that the cell-to-cell propagation of virus infection is linked to the ability of the viral MP to suppress the dsRNA-induced defense response leading to PD closure. Taken together, the results draw a central role of PTI signaling and suppression in determining the ability of viruses to spread infection between cells in susceptible plants.

## Results

### dsRNA causes inhibition of virus movement in *N. benthamiana*

To discover how dsRNA-induced PTI inhibits RNA virus infection, we visualized the effect of poly(I:C) treatment on local infections of *Nicotiana benthamiana* plants using tobacco mosaic virus tagged with green fluorescent protein (TMV-GFP). The TMV-GFP infection sites were lower in number and smaller at 7 days post inoculation (dpi) in plants treated with poly(I:C) or with a bacterial PTI elicitor derived from flagellin (flg22) than in control plants treated with water (**Figure 1, A** and **B**, and **Figure S1**). The treatments did not cause a significant change in GFP fluorescence intensity, an indicative for viral replication and accumulation (**Figure 1A**) indicating that they may not exert a significant bulk effect on viral RNA accumulation in infected cells. To test this further, we measured the accumulation of viral RNA in leaves agroinfiltrated for expression a cell-autonomous, MP-deficient TMV replicon (TMVΔMΔC-GFP). As is shown in **Figure 1C**, pre-treatment of the leaves with poly(I:C) did not elicit a significant effect on TMVΔMΔC-GFP viral accumulation through a time-course of infection at 1, 3, and 5 days post infection (dpi) compared to leaves treated with water. Therefore, the reduced size and number of infection sites in poly(I:C)-treated leaves suggested that the poly(I:C)-triggered immunity may be linked to the reduced cell-to-cell movement of the virus.

**Figure 1.**
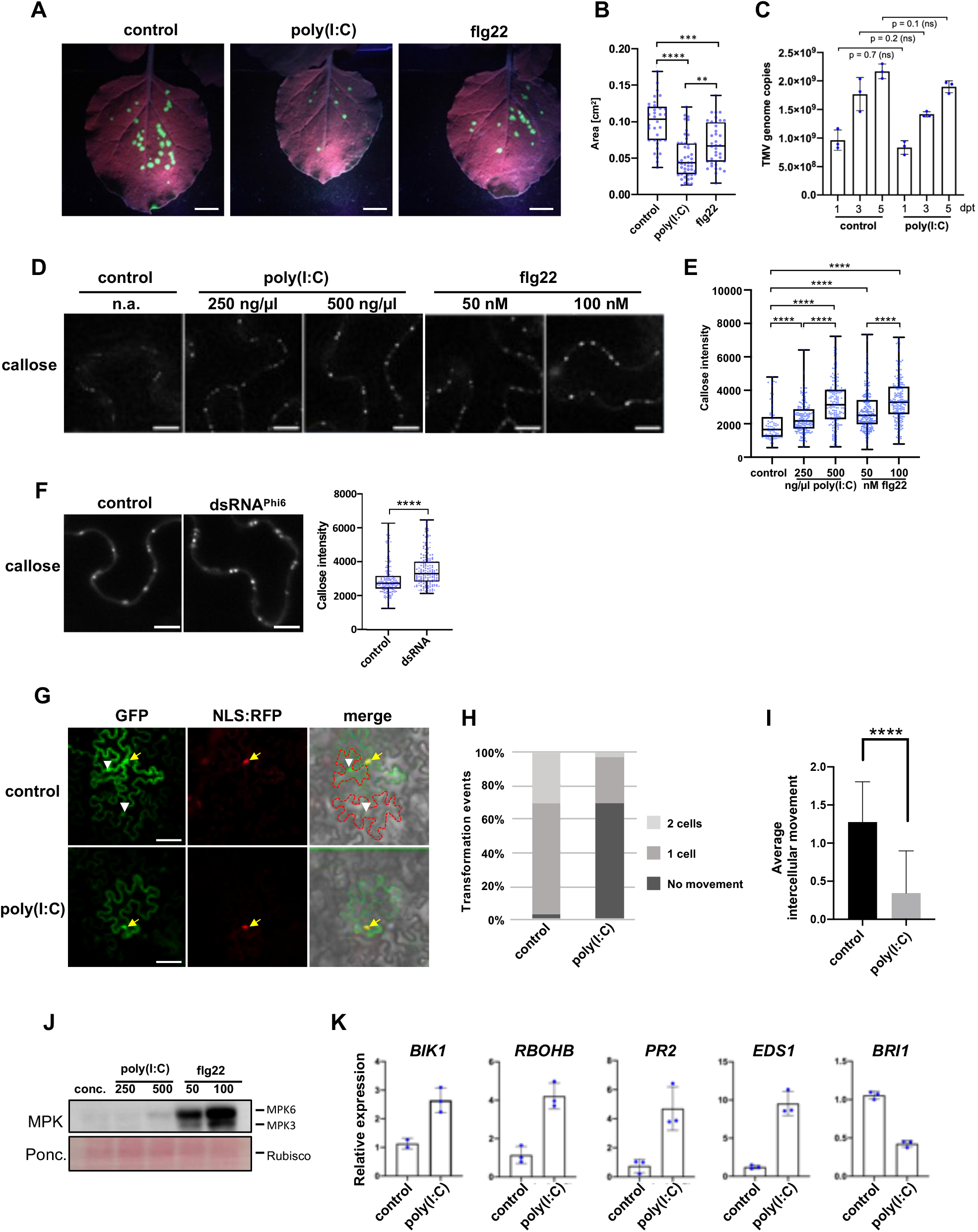
Poly(I:C) treatment causes inhibition of virus movement in *N. benthamiana*. (**A**) TMV-GFP infection sites in *N. benthamiana* leaves at 7 days post inoculation (dpi) with the virus together with either water (control), 0.5 μg/μl (≈1 μM) poly(I:C), or 1 μM flg22. Scale bar, 1 cm. (**B**) Sizes of individual infection sites measured in 10 leaf samples collected from three plants per treatment. Two-tailed Mann-Whitney test; ****, p <0.0001; ***, p <0.001; **, p <0.01. The experiment was performed three times with similar results. (**C**) TMV replication in *N. benthamiana* is not influenced by poly(I:C). A cell-autonomous, MP-deficient TMV replicon (TMVΔMΔC-GFP) expressed in cells of agroinoculated leaves produces the same number of RNA genome copies in the presence and absence of treatment with 0.5 μM poly(I:C), as determined by Taqman RT-qPCR. Poly(I:C) and control treatments were applied 1 day after agroinoculation and results obtained at indicated days after this treatment (dpt) are shown. The data represent means of three biological replicates (with SD) per time point and treatment. Two-tailed Mann-Whitney test. ns, not significant. (**D**) and (**E**) Treatment of *N. benthamiana* with poly(I:C) or flg22 induces increased callose deposition at PD within 30 minutes in a dose-specific manner. (**D**) Callose fluorescence at PD upon aniline blue staining. Scale bar, 10 μm. (**E**) Relative callose content in individual PD (blue dots, n > 100) as determined in three leaf discs per treatment. Two-tailed Mann-Whitney test; ****, p = <0.0001. (**F**) Callose deposition at PD in *N. benthamiana* leaf epidermal tissue upon treatment with 50 ng/μl biological dsRNA (dsRNA^Phi6^). Relative callose content in individual PD (blue dots, n > 100) as determined in three leaf discs per treatment. Two-tailed Mann-Whitney test; ****, p = <0.0001. (**G-I)** GFP mobility assay in *N.* *benthamiana*. Leaf disks expressing GFP together with cell-autonomous NLS:RFP one day after agroinfiltration were treated with water or 0.5 μg/μl poly(I:C) and imaged 48 hours later. (**G)** Example of GFP movement from an epidermal cell marked by cell-autonomous NLS:RFP into adjacent cells. Transiently expressed GFP shows a nucleocytoplasmic distribution (yellow arrow) and its movement from the expressing epidermal cell (co-expressed NLS:RFP in the nucleus, in red) is evident by appearance of green fluorescence in the nuclei and cytoplasm of adjacent cells (white arrowheads). Cells into which GFP moved are indicated by the red dashed line in the merged image. Scale bar, 50 μm. (**H** and **I)** Quantification of GFP movement between epidermal cells in leaf disks exposed to 0.5 μg/μl poly(I:C) or water (control) (29 transformation events were analyzed for each treatment). (**H**) Stacked column diagram showing the relative frequency of transformation events associated with either no GFP movement (dark grey), GFP movement into one adjacent cell layer (medium grey), or GFP movement into two adjacent cell layers (light grey). (**I**) Average intercellular movement (total number of cell layers into which GFP has moved divided by the number of evaluated transformation events). Two-tailed Mann-Whitney test; ****, p = <0.0001. A repetition of the GFP mobility assay provided similar results. (**J**) Low level of MPK activation by poly(I:C) relative to flg22 after 30 minutes. Concentrations (conc.) are in ng/μl for poly(I:C) and in nM for flg22. The experiment was performed three times with similar results. (**K**) Poly(I:C) induces innate immunity marker genes, but suppresses expression of *BRI1*, in *N. benthamiana*. Mean value and SD of gene expression values obtained by RT-qPCR with three biological replicates (blue dots) harvested three hours after treatment.

### dsRNA triggers callose deposition at PD along with the activation of typical PTI responses in *N. benthamiana*

Because virus intercellular movement occurs through PD (Heinlein, 2015) we hypothesized that dsRNA inhibits virus movement by causing PD closure. A major mechanism restricting the conductivity of PD for the transport of macromolecules involves the deposition of callose (β-1,3-glucan) in the cell wall region surrounding the PD channel (Wu et al., 2018). Consistently, treatment of *N. benthamiana* plants with poly(I:C), flg22, or water, and quantification of PD-associated callose by *in vivo* aniline blue staining (Huang et al., 2022) revealed that both poly(I:C) and flg22 trigger increased levels of PD-associated callose in a concentration-dependent manner (**Figure 1, D** and **E**). A similar induction of callose at PD is also seen upon treatment of *N. benthamiana* plants with Phi6 dsRNA (Niehl et al., 2018) (**Figure 1F**), which dismisses the possibility that poly(I:C) induced callose deposition through an unspecific effect. In agreement with poly(I:C)-induced callose deposition at PD, poly(I:C)-treated tissues showed reduced PD permeability as determined by a GFP mobility assay. In this assay, isolated individual cells of *N. benthamiana* leaves were transformed for the expression of cytoplasmic GFP together with red fluorescent protein tagged with a nuclear localization signal (NLS-RFP) as a red fluorescent cell-autonomous marker. Whereas more than 97% of the observed GFP-expressing cells showed GFP mobility into one or two adjacent cell layers in control (water)-treated tissues, this mobility was reduced to 31% in the presence of poly(I:C) (**Figure 1, G - I**). Moreover, as previously noted in Arabidopsis (Niehl et al., 2016), poly(I:C) triggered a moderate MPK activation and the level of activation is significantly weaker than the activation observed with flg22 (**Figure 1J**). Poly(I:C)-treated leaves also exhibited the induction of *N. benthamiana* defense-related genes, such as genes encoding BOTRYTIS INDUCED KINASE1 (BIK1), PATHOGENESIS-RELATED PROTEIN 2 (PR2), NADPH/RESPIRATORY BURST OXIDASE PROTEIN B (RBOHB) and ENHANCED DISEASE SUSCEPTIBILITY 1 (EDS1), whereas the gene for BRASSINOSTEROID INSENSITIVE 1 (BRI1) was down-regulated (**Figure 1K**).

### Poly(I:C)-induced PD callose deposition in Arabidopsis requires PTI signaling components but is independent of ROS and MPK

To determine how dsRNA elicits the deposition of callose at PD, we turned our attention to Arabidopsis. As noted previously (Niehl et al., 2016) the treatment of *A. thaliana* Col-0 plants with poly(I:C) causes a significant induction in some PTI-related gene expression, including *SERK1* (**Figure 2A**), whereby the induction of *PR5* was found to be SERK1-dependent (**Figure 2B**). Moreover, the same treatment as well as the treatment with 50 ng/μl phi6 dsRNA increased the PD-associated callose levels as seen before in *N. benthamiana* (**Figure 2, C and D**). The induced callose depositions are exactly localized to PD as shown in transgenic *A. thaliana* Col-0 plants expressing PD markers PLASMODESMATA CALLOSE BINDING 1 fused to the red fluorescent protein mCHERRY (mCherry-PDCB1) or the Plasmodesmata-localized β-1,3 glucanase 2 fused to the yellow fluorescent protein mCitrine (PdBG2-mCitrine) (Benitez-Alfonso et al., 2013) (**Figure 2, E** and **F**). Interestingly, dsRNA-induced callose deposition was strongly inhibited in *bik1 pbl1* plants (**Figure 2G**), which are deficient in the receptor-like cytoplasmic kinase (RLCK) BIK1 and its homolog PBS1-LIKE KINASE1 (PBL1). The BIK1 receptor-like cytoplasmic kinase (RLCK) module is an important component of PTI signaling that integrates signals from multiple pathogen-recognition receptors (PRRs), as shown by its direct interaction with the PRR proteins FLAGELLIN SENSITIVE 2 (FLS2), EF-TU RECEPTOR (EFR), PEPTIDE RECEPTOR (PEPR)1 and PEPR2, and CHITIN ELICITOR RECEPTOR KINASE 1 (CERK1) (Lu et al., 2010; Zhang et al., 2010; Liu et al., 2013), and its ability to phosphorylate and activate downstream targets, such as the NADPH Oxidase RESPIRATORY BURST OXIDASE HOMOLOG D (RBOHD) (Kadota et al., 2014). BIK1 and PBL1 have additive effects; unlike the single mutants, the *bik1 pbl1* double mutant was shown to strongly inhibit PAMP-induced defense responses (Zhang et al., 2010). In addition, *bik1 pbl1* plants are also deficient in poly(I:C)-induced MPK activation and seedling root growth inhibition as compared to Col-0 wild type (WT) plants (**Figure 2, H and I**).

**Figure 2.**
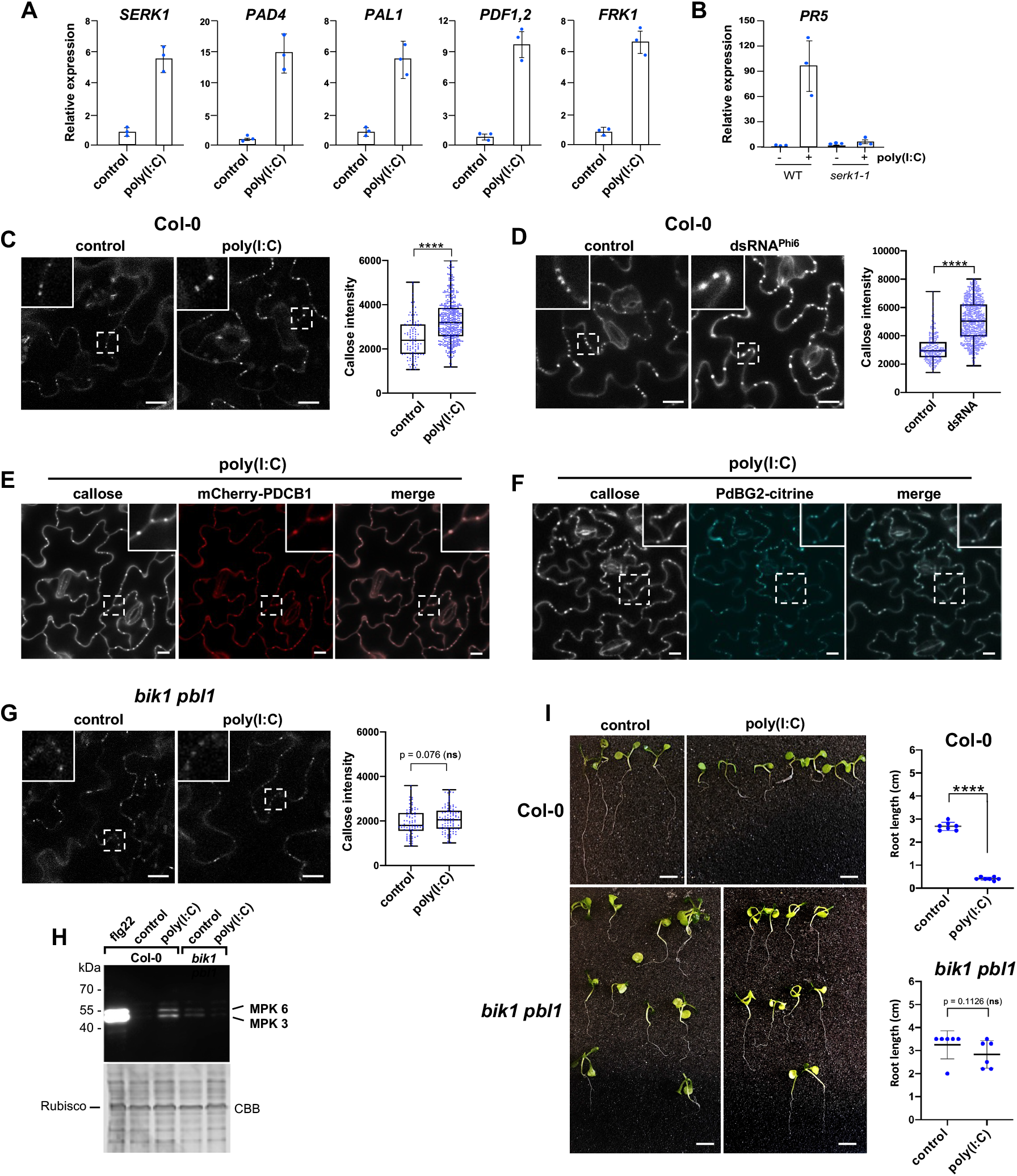
Poly(I:C)-induced signaling in Arabidopsis depends on BIK1/PBL1. **(A)** and **(B)** Transcriptional regulation of Arabidopsis genes three hours after treatment with 0.5 μg/μl poly(I:C). For each gene, the mean value and the SD of gene expression values obtained by RT-qPCR analysis of three biological replicates (blue dots) is shown. (**A**) Poly(I:C) induces innate immunity marker genes in *A. thaliana* Col-0 wildtype. (**B**) Absence of poly(I:C)-induced *PR5* expression in the *serk1-1* mutant. **(C**) Poly(I:C) treatment causes callose deposition at PD in *A. thaliana* Col-0. Images were taken 30 minutes after treatment. Inlays show enlargements of the areas within the dashed boxes. Scale bar, 20 μm. Relative callose content in individual PD (blue dots, n > 100) as determined in three leaf discs from three plants per treatment. Two-tailed Mann-Whitney test; ****, p = <0.0001. (**D**) Callose deposition at PD in *A. thaliana* Col-0 leaf epidermal tissue 30 minutes after treatment with 50 ng/μl of biological dsRNA (dsRNA^Phi6^). Inlays show enlargements of the areas within the dashed boxes. Scale bar, 20 μm. Relative callose content in individual PD (blue dots, n > 100) as determined in three leaf discs per treatment. Two-tailed Mann-Whitney test; ****, p = <0.0001. (**E**) and (**F**) Poly(I:C)-induced callose spots are localized to PD as shown by co-localization with PD markers mCherry-PDCB1 (**E**) and PdBG2-citrine (**F**). Inlays show enlargements of the areas within the dashed boxes. Scale bar, 10 μm. (**G**) poly(I:C)-induced callose deposition at PD is inhibited in the *bik1 pbl1* mutant Images were taken 30 minutes after treatment and the WT control of the same experiments is shown in (**C**). Inlays show enlargements of the areas within the dashed boxes. Scale bar, 20 μm. Relative callose content in individual PD (blue dots) as determined in three leaf discs per treatment. Two-tailed Mann-Whitney test; ns, non-significant. (**H**) Poly(I:C)-induced MPK activation is reduced in the *bik1 pbl1* mutant. Immunoblot detection of phosphorylated MPK. Samples were harvested 30 minutes after treatment with 0.5 μg/μl poly(I:C), 1 μM flg22, or water. “*bik1*” stands for *bik1 pbl1*. CBB, Coomassie brilliant blue-stained gel showing staining of ribulose-bisphosphate-carboxylase (Rubisco) as gel loading control. (**I**) *bik1 pbl1* plants do not show significant seedling root growth inhibition in the presence of poly(I:C) as compared to WT Col-0 plants. Seedlings were kept for 12 days in 0.5 μg/μl poly(I:C) or water. Scale bar, 1 cm. Quantification of poly(I:C)-induced root growth inhibition in *A. thaliana* WT Col-0 and *bik1 pbl1* seedlings. Analysis of 6-7 seedlings (blue dots) per condition. Two-tailed Mann-Whitney test; ****, p < 0.0001; ns, non-significant.

Perception of flg22 by the FLS2 and BAK1 (BRASSINOSTEROID INSENSITIVE1 (BRI1)-ASSOCIATED RECEPTOR KINASE1) co-receptor complex induces rapid phosphorylation of BIK1, evidenced by a protein mobility shift in immunoblotting analysis (Lu et al., 2010; Zhang et al., 2010). To determine if BIK1 is phosphorylated in the presence of poly(I:C), Arabidopsis WT Col-0 protoplasts expressing HA epitope-tagged BIK1 were treated with poly(I:C) for 20 minutes. Subsequent Western blot analysis with anti HA-antibody revealed that similar to flg22 treatment, poly(I:C) treatment induces a mobility shift of BIK1-HA proteins (**Figure 3A**). BIK1 phosphorylation was confirmed by the absence of this mobility shift in the presence of calf intestine phosphatase (CIP) (**Figure 3, A and B**) or the protein kinase inhibitor K-252a (**Figure 3B**). Taken together, the data suggest the involvement of BIK1/PBL1 in dsRNA-triggered immunity mediating the callose deposition at the PD.

**Figure 3.**
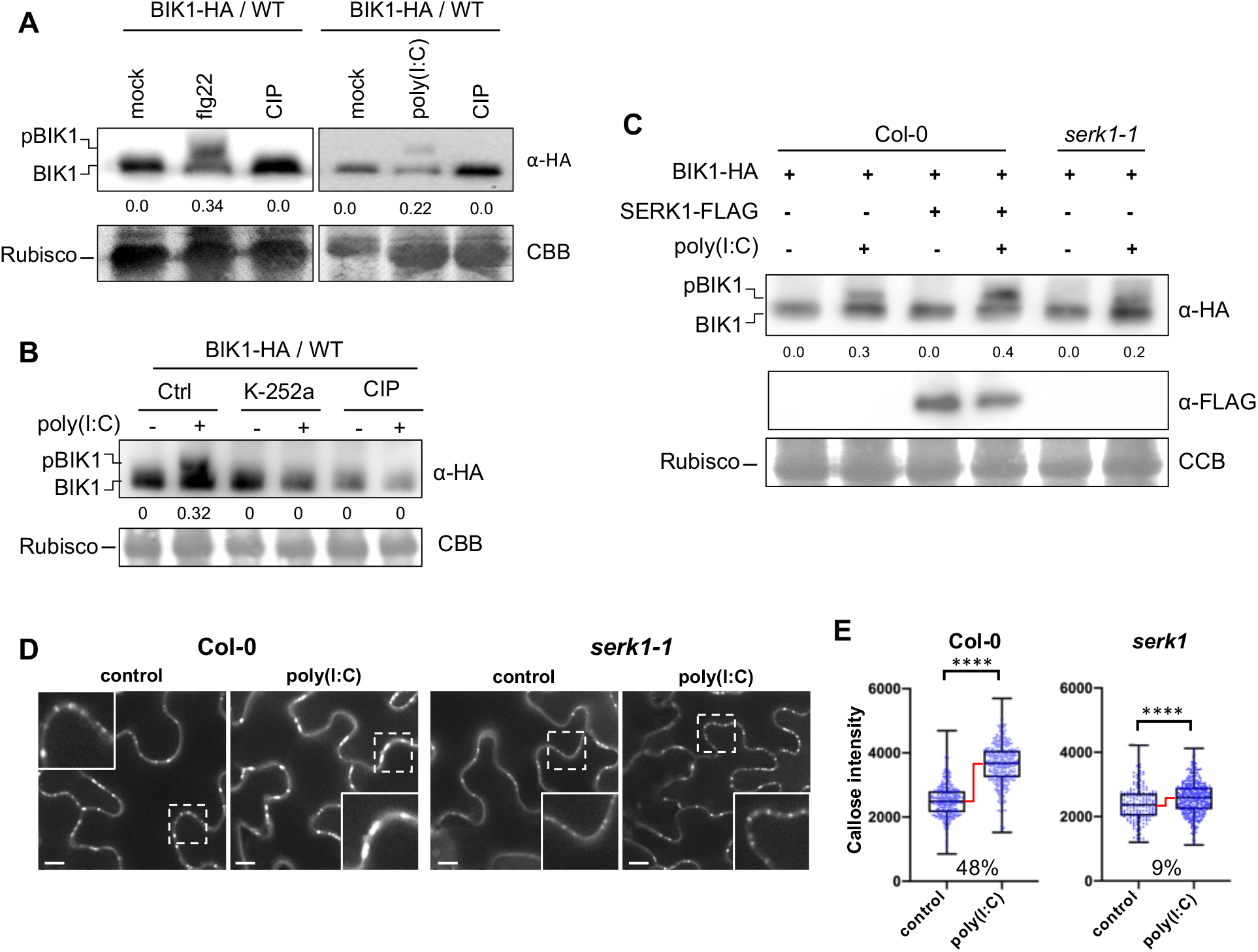
Poly(I:C) causes BIK1 phosphorylation and PD callose deposition in a SERK1-dependent manner. (**A**) to (**C**) Analysis if BIK1 phosphorylation in poly(I:C)-treated *A. thaliana* Col-0 protoplasts. (**A**) Poly(I:C) treatment induces BIK1 phosphorylation as shown by a protein mobility shift detected by Western blot analysis. Protoplasts expressing BIK1-HA were non-treated (mock) or treated with 1 μM flg22 or 0.5 μg/μl poly(I:C), lysed after 20 minutes and treated or non-treated with calf intestine phosphatase (CIP) for 60 before Western blot analysis using HA-HRP antibody. BIK1 band intensities were quantified using Image Lab (Bio-Rad). Quantification of BIK1 phosphorylation (upper panel) calculated as ratio of intensity of the upper band (phosphorylated BIK1, pBIK1) to the sum intensities of shifted and non-shifted bands (pBIK1 + BIK1) (no treatment set to 0.0). CCB, Coomassie brilliant blue staining of Rubisco as gel loading control (lower panel). (**B**) Poly(I:C)-induced BIK1 phosphorylation is blocked by 1 μM of the kinase inhibitor K-252a added 1 hour before poly(I:C) treatment. Rubisco detection by CBB staining is shown as gel loading control (lower panel). Experimental conditions and quantification of BIK1 phosphorylation as in (**A**). (**C**) SERK1 enhances poly(I:C)-induced BIK1 phosphorylation. Protoplasts from WT Col-0 or *serk1-1* mutants were transfected with BIK1-HA together with or without SERK1-FLAG and followed by treatment with or without 0.5 μg/μl poly(I:C). Phosphorylated BIK1 band intensities were quantified as in (**A**). The middle panel shows SERK1-FLAG expression. Rubisco detection by CBB staining is shown as gel loading control (lower panel). (**D**) and (**E**) Poly(I:C)-induced PD callose deposition depends on SERK1. (**D**) Callose fluorescence at PD seen upon aniline blue staining of epidermal cells of WT Col-0 plants and *serk1-1* mutants treated with water (control) or 0.5 μg/μl poly(I:C). Inlays show enlargements of the areas within the dashed boxes. Scale bar, 10 μm. (**E**) Relative callose content (fluorescence intensity) in individual PD (blue dots, n > 100) as determined in three leaf discs taken from three plants per treatment. Two-tailed Mann-Whitney test; ****, p = < 0.0001. The increase in PD callose levels in poly(I:C)-treated samples relative to water control-treated samples is shown in percent (%). Although the callose intensity data distributions between poly(I:C)-treated and control-treated samples are significantly different in both WT Col-0 and *serk1*, the comparison of the callose intensity median levels indicate a drastic inhibition of the poly(I:C)-induced PD callose deposition response in *serk1* as compared to the WT.

Previously, we showed that *serk1-1* mutants show the reduced levels of poly(I:C)-induced ethylene production and antiviral protection [Niehl, 2016 #6670]. We further investigated the involvement of SERK1 in poly(I:C)-induced BIK1 phosphorylation. As is shown in **Figure 3C**, the level of poly(I:C)-induced BIK1 phosphorylation was increased upon SERK1 overexpression in WT Col-0 plants and decreased in the *serk1-1* mutant. Importantly, the poly(I:C)-induced PD callose deposition was drastically reduced in *serk1-1* mutants compared to WT Col-0 plants (**Figure 3D** and **E**). Whereas the median PD callose level increased by 48% in WT col-0 plants upon poly(I:C) treatment, only a minor increase in the median PD callose level was observed in the *serk1* mutant. Thus, the data indicate that SERK1 contributes to poly(I:C)-induced BIK1 phosphorylation and may function genetically upstream of BIK1.

To investigate the poly(I:C)-induced signaling pathway downstream of BIK1, we examined the production of reactive oxygen species (ROS) by a luminescence assay. ROS play important roles in plant development and stress responses (Mittler, 2017) and are also produced during infections with fungal and bacterial pathogens (Castro et al., 2021). ROS accumulate also upon perception of the fungal and bacterial elicitors chitin and flagellin (flg22) (Nuhse et al., 2007; Cheval et al., 2020) and have been linked to local and systemic signaling, including calcium signaling, and the deposition of callose at PD (Faulkner et al., 2013; Cheval et al., 2020). Notably, neither the treatment of Arabidopsis Col-0 plants (**Figure 4A**) nor the treatment of *N. benthamiana* plants (**Figure 4B**) with poly(I:C) led to the production of ROS. By contrast, strong responses were recorded in both plant species upon treatment with the flg22 elicitor. In addition, *rbohd* and *rbohf* mutants deficient for the major ROS-producing NADPH oxidases RESPIRATORY BURST OXIDASE HOMOLOG D (RBOHD) and RBOHF (Castro et al., 2021) responded like WT Col-0 plants to the presence of poly(I:C) in showing induced callose deposition at PD (**Figure 4C**). Thus, unlike for chitin and flagellin (flg22), the induction of PD callose deposition by poly(I:C) is likely independent of ROS. Moreover, *mpk3* and *mpk6* single mutants that are deficient for the mitogen activated protein kinases (MPK) 3 and 6, respectively, as well as the *mpk3 amiRmpk6* mutants (an *mpk3* mutant in which *MPK6* is silenced by an artificial miRNA) (Li et al., 2014) showed increased levels of callose at PD upon poly(I:C) treatment similar to WT plants (**Figure 4D**). Considering the relatively weak activation of MPKs by poly(I:C) treatment (**Figure 1J** and **Figure 2H**), it is possible that the MPK3/6 module may not play a major role in dsRNA-induced callose deposition. Alternatively, it is also possible that additional yet non-identified MPKs may be involved in this process.

**Figure 4.**
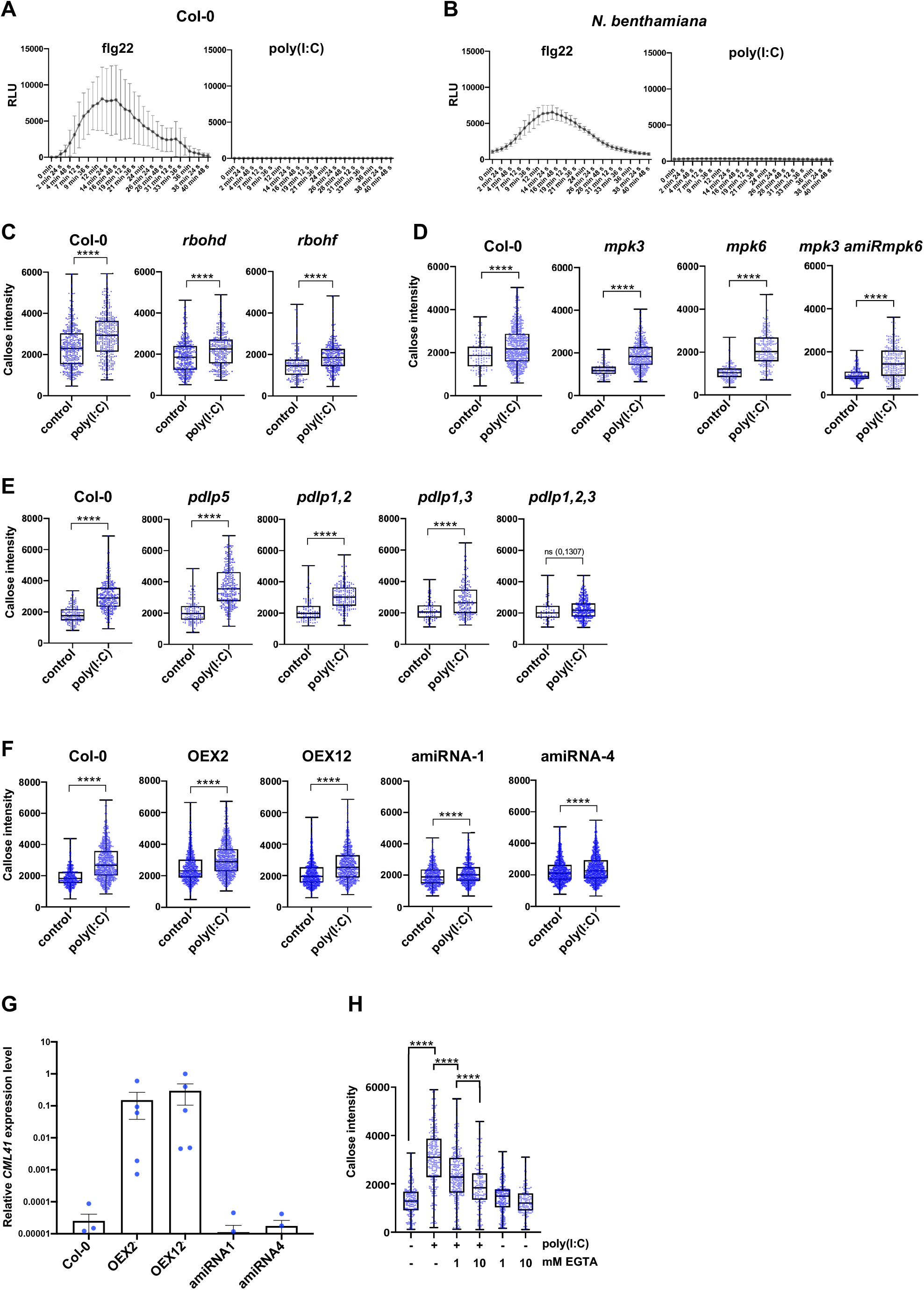
poly(I:C)-induced PD callose deposition is ROS- and MPK3/6 cascade-independent but requires PDLP1/2/3 and CML41. (**A**) and (**B**) Unlike flg22, poly(I:C) treatment does not induce any ROS production in Arabidopsis (**A**) or *N. benthamiana* (**B**). RLU, relative luminescence units. Mean values (black dots) and error bars (SD) obtained for each time point for 10 replicates (leaf disks) per treatment. (**C**) Poly(I:C)-induced callose deposition at PD is not affected in *rbohd* or *rbohf* mutants. Relative callose content in individual PD (blue dots, n > 100) 30 minutes after treatment with 0.5 μg/μl poly(I:C) or water and determined in three leaf discs from three plants per treatment. Two-tailed Mann-Whitney test; ****, p = <0.0001. (**D**) Poly(I:C)-induced callose deposition at PD is not affected in *mpk3* and *mpk6* single mutants, and neither in a *mpk3* mutant in which *MPK6* is silenced by an artificial miRNA (*mpk3 amiRmpk6*). Relative callose content in individual PD (blue dots, n > 100) 30 minutes after treatment with 0.5 μg/μl poly(I:C) or water and determined in three leaf discs from three plants per treatment. Two-tailed Mann-Whitney test; ****, p = <0.0001. (**E**) Poly(I:C)-induced callose deposition at PD is independent of PDLP5 but depends on the redundantly acting PDLP1, PDLP2, and PDLP3. Relative callose content in individual PD (blue dots, n > 100) 30 minutes after treatment with 0.5 μg/μl poly(I:C) or water and determined in three leaf discs from three plants per treatment. Two-tailed Mann-Whitney test; ****, p = <0.0001; ns, non-significant. (**F**) and **(G**) Poly(I:C)-induced callose deposition at PD depends on CML41. (**F**) PD callose deposition levels in poly(I:C)-treated and control-treated leaf disks of two *CML41*-overexpressing lines (OEX2 and OEX12) and of two lines in which the expression of *CML41* is reduced by expression of artificial miRNA (amiRNA1 and amiRNA4). As compared to the WT (Col-0) and the *CML41*-overexpressing lines, the inducibility of PD callose deposition by poly(I:C) is strongly decreased in the amiRNA lines. Relative callose content in individual PD (blue dots, n > 100) 30 minutes after treatment with 0.5 μg/μl poly(I:C) or water and determined in three leaf discs from three plants per treatment. Two-tailed Mann-Whitney test; ****, p = <0.0001. According to the Mann-Whitney test the distribution of individual PD callose intensities is different between all control- and poly(I:C)-treated samples. However, unlike in WT Col-0 and *CML41*-overexpressing lines, the median PD callose intensity levels are not increased upon poly(I:C)-treatment in amiRNA lines thus indicating a deficiency in the induction of PD callose deposition upon poly(I:C) treatment in these lines. (**G**) Relative levels of *CML41* expression in plants of the OEX2, OEX12, amiRNA1 and amiRNA4 lines in comparison to WT (Col-0), as determined by RT-qPCR. Mean value and standard error of gene expression values obtained by RT-qPCR with 3-6 biological replicates (blue dots). (**H**) poly(I:C)-induced callose deposition is reduced in the presence of EGTA. Relative callose content in individual PD (blue dots, n > 100) as determined in three leaf discs per treatment. Two-tailed Mann-Whitney test; ****, p = <0.0001.

To further investigate the signaling pathway induced by dsRNA, additional mutants were tested. We started with Arabidopsis mutants deficient in the PD-localized proteins (PDLPs), which are a family of eight proteins that dynamically regulate PD (Thomas et al., 2008). PDLP5 plays a non-redundant role in intercellular systemic acquired resistance (SAR) signaling (Lim et al., 2016) and in mediating salicylic acid (SA)-induced PD closure, a process required for resistance against the bacterial pathogen *P. syringae* (Lee et al., 2011; Wang et al., 2013). However, *pdlp5* mutant plants showed strong callose deposition at PD upon poly(I:C) treatment (**Figure 4E**), indicating that dsRNA-induced callose deposition is independent of PDLP5 and of a potential SA response mediated by this protein. Next, we tested a PDLP1, PDLP2 and PDLP3, which play redundant roles in callose deposition at PD (Thomas et al., 2008), in callose deposition within haustoria formed in response to infection by mildew fungus (Caillaud et al., 2014), and also as binding receptors for tubule-forming viruses (Amari et al., 2010). Whereas *pdlp1 pdlp2* (*pdlp1,2*) and *pdlp1 pdlp3* (*pdlp1,3*) double mutants showed a normal poly(I:C)-induced callose deposition, *pdlp1 pdlp2 pdlp3* (*pdlp1,2,3*) triple mutant plants were unable to significantly increase PD-associated callose levels in response to poly(I:C) (**Figure 4E**). This observation suggests the involvement of PDLP1, PDLP2 and PDLP3 in dsRNA-triggered immunity and in mediating the callose deposition at the PD.

In further screening of other mutants for dsRNA sensitivity, we found that poly(I:C)-induced callose deposition at PD also depends on the Ca^2+^-binding, PD-localized CALMODULIN-LIKE protein 41 (CML41). This protein was shown to mediate rapid callose deposition at PD associated with a decreased PD permeability following flg22 treatment (Xu et al., 2017). Plants of *CML41* overexpressing transgenic lines (*CML41-OEX-2* and *CML41-OEX-12*) (Xu et al., 2017) showed increased PD-associated callose levels upon poly(I:C) treatment similar to WT Col-0 plants (**Figure 4F**). In contrast, transgenic plant lines in which *CML41* is downregulated by an artificial miRNA (*CML41-amiRNA-1* and *CML41-amiRNA-4*) (Xu et al., 2017) showed a seven- to eight-fold lower ability to respond to this treatment as compared to WT Col-0 (**Figure 4F**). The reduction in the response to poly(I:C) of *CML41-amiRNA* plants correlates with the reduced level of *CML41* expression in these lines (Xu et al., 2017) (**Figure 4G**). Consistent with the role of CML41 in the callose deposition response to poly(I:C), the permeability of PD was previously shown to be sensitive to cytosolic Ca^2+^ concentrations (Tucker and Boss, 1996; Holdaway-Clarke et al., 2000). To test the role of Ca^2+^ in dsRNA-triggered innate immunity, we treated plants with poly(I:C) together with EGTA, a Ca^2+^-chelating molecule. The level of callose induced at PD after dsRNA treatment was reduced in the presence of EGTA in a concentration-dependent manner (**Figure 4H**), indicating a role of Ca^2+^ in poly(I:C)-triggered PD regulation. Together, these results suggest a role for CML41 and Ca^2+^ in the poly(I:C)-induced defense response at PD. The observation that the poly(I:C)-induced callose levels were not significantly increased by *CML41* overexpression may be due to that the endogenous level of *CML41* expression in WT plants is sufficient for the full activity of the PD-localized CML41 proteins. However, the level of expression in WT plants is critical as a reduction in *CML41* levels strongly affected PD callose levels.

### BIK1/PBL1 and CML41 are essential for dsRNA-induced antiviral resistance

Previously, we showed that poly(I:C) co-treatment during virus inoculation protects Arabidopsis plants against infection by oilseed rape mosaic virus (ORMV) and that efficient protection depends on SERK1 (Niehl et al., 2016). As we demonstrate here, SERK1 plays an essential role in the poly(I:C)-induced callose deposition at PD, implying a role of PD closure in dsRNA-induced antiviral resistance. To further test the significance of PD callose deposition of BIK1/PBL1 and CML41 in dsRNA-induced antiviral resistance, we inoculated poly(I:C)-treated and non-treated *bik1 pbl1* and *CML41 amiRNA-1* plants with ORMV. Whereas poly(I:C) treatment prevented symptoms at 28 dpi and resulted in a strongly reduced virus titer in WT Col-0 plants, *bik1 pbl1* and *CML41 amiRNA-1* plants showed severe virus-infected symptoms and accumulated high virus levels in poly(I:C)-treated plants similar to those plants without poly(I:C) treatment upon ORMV infection (**Figure 5, A** and **B**). Ablation of virus-inoculated leaves from plants at different times after inoculation showed that the time required for the virus to exit the inoculated leaf and to cause systemic infection was three days in WT plants. By contrast, this time was reduced to 24 hours in *bik1 pbl1* mutants and *CMLl41-amiRNA-1* plants (**Figure 5, C** and **D**). These findings show that dsRNA-induced antiviral PTI occurs at the level of virus movement. Consistent with this PTI effect on virus movement, the experiments reveal a dsRNA-induced signaling pathway that requires SERK1, BIK1/PBL1, CML41, Ca^2+^ and PDLP1/2/3 for callose deposition at PD. This dsRNA-induced callose deposition at PD is likely independent of ROS and MPK3/6 signaling, which differs from the immune signaling triggered by fungal and bacterial elicitors (Kadota et al., 2014; Cheval et al., 2020).

**Figure 5.**
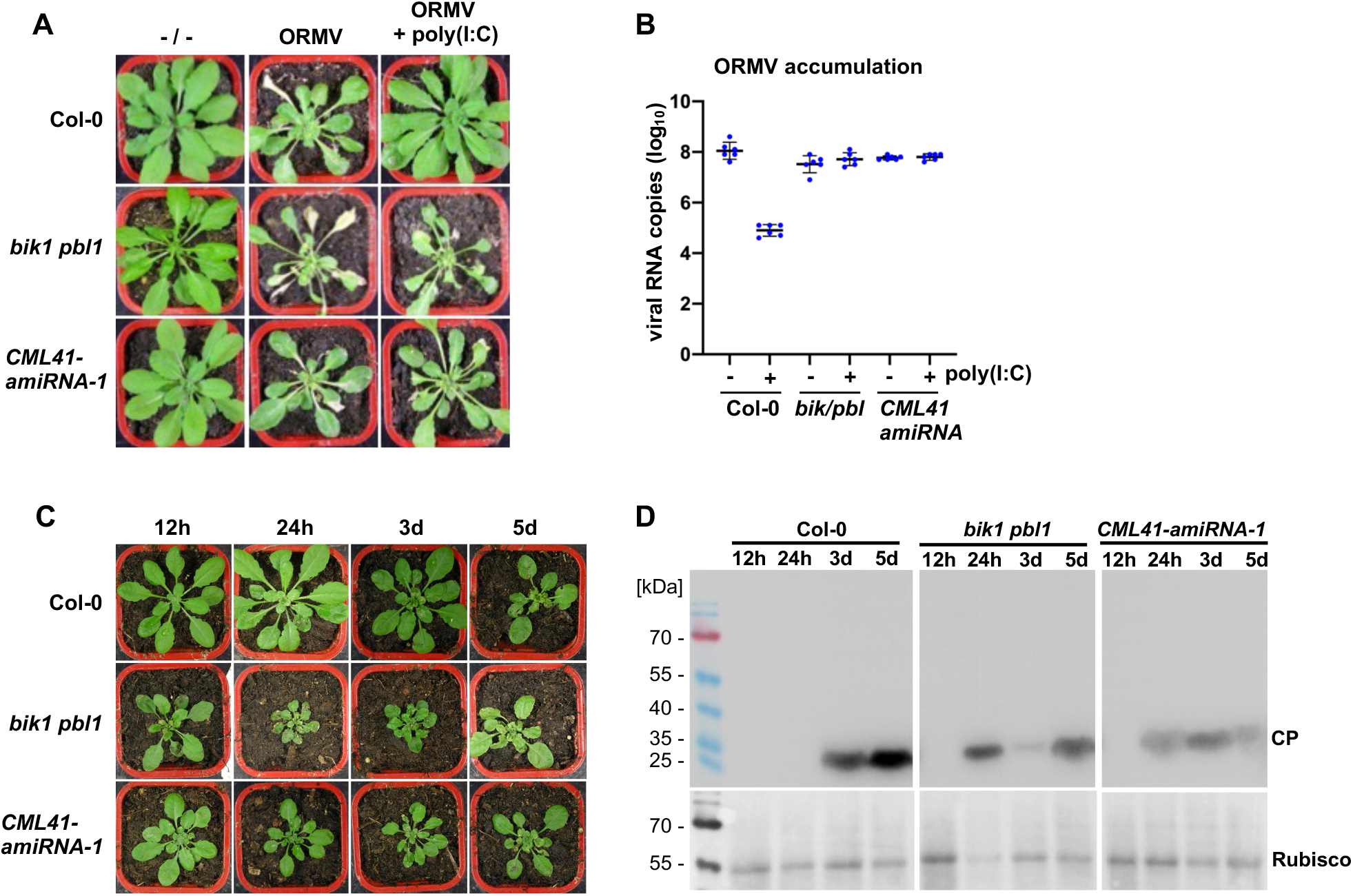
BIK1/PBL1 and CML41 are required for antiviral defense. (**A**) and (**B**) Disease symptoms (**A**) and viral RNA accumulation (**B**) at 28 dpi in wild-type plants and mutants inoculated with ORMV in the presence and absence of 0.5 μg/μl poly(I:C). Unlike in wild-type plants (Col-0) the antiviral effect of poly(I:C) treatment is lost in *bik1 pbl1* mutants and *CML41-amiRNA-1* expressing plants. Viral RNA accumulation (**B**) is depicted for 6 biological replicates per condition. Mean values and standard errors are shown. (**C**) and (**D**) BIK1 and CML41 inhibit virus movement. (**C**) Representative symptom phenotypes at 21 dpi of Arabidopsis Col-0 plants, *bik1 pbl1* mutant plants and plants transgenic for *CML41-amiRNA-1* that were locally inoculated with ORMV and from which the inoculated leaves were removed at the indicated times in hours (h) and days (d). Whereas systemic leaves of Col-0 plants show symptoms on plants that carried the inoculated leaves for 3 or more days following inoculation, the systemic leaves of the *bik1 pbl1* mutant and of the CML41-amiRNA-expressing plants show symptoms already if the inoculated leaves were present for only 24 hours. (**D**) Immunoblot analysis of the youngest systemic leaves at 21 dpi using antibodies against viral coat protein (CP) (Youcai mosaic virus antibody, AS-0527, DSMZ, Braunschweig, Germany). The pattern of CP expression in the systemic leaves confirms that in WT Col-0 plants the virus needs between 24 h and 3 d to exit the inoculated leaves and move systemically, whereas the time needed for systemic movement is reduced to less than 24 h in the *bik1 pbl1* mutant and of the CML41-amiRNA expressing plants, thus indicating a role of BIK1 and CML41 in restricting virus movement.

### dsRNA-induced callose deposition is suppressed by viral movement protein

The plant-pathogen arms race causes pathogens to evolve virulent effectors that overcome host defenses. Viral MPs are essential in mediating virus movement during infection. We tested whether viral MPs are involved in the suppression of the dsRNA-induced callose deposition at PD. To address this question, we divided the local TMV infection site into different zones (**Figure 6A**): zone I ahead of infection and without MP, zone II at the virus front where MP facilitates virus movement, zone III behind the infection front, and zone IV, which is the center of the infection site where MP is no longer expressed. *In vivo* detection of dsRNA with GFP-fused dsRNA-binding protein B2 of *Flock house virus* (Monsion et al., 2018) shows that zones II-IV accumulate dsRNA in distinct replication complexes that also produce MP (**Figure 6B**). Aniline blue staining demonstrates high PD-associated callose levels within and around the infection site (**Figure 6C**). However, cells in zone II and zone III, where virus cell-to-cell movement is associated with a transient activity of MP in increasing the PD size exclusion limit (SEL) (Oparka et al., 1997), exhibit a marked reduction in PD-associated callose levels as compared to cells in zone I (ahead of infection) and zone IV (center of infection) (**Figure 6, C** and **D**). The low level of PD-associated callose in cells at the virus front (zone II) is consistent with the ability of MP to interfere with dsRNA-triggered immunity leading to PD closure.

**Figure 6.**
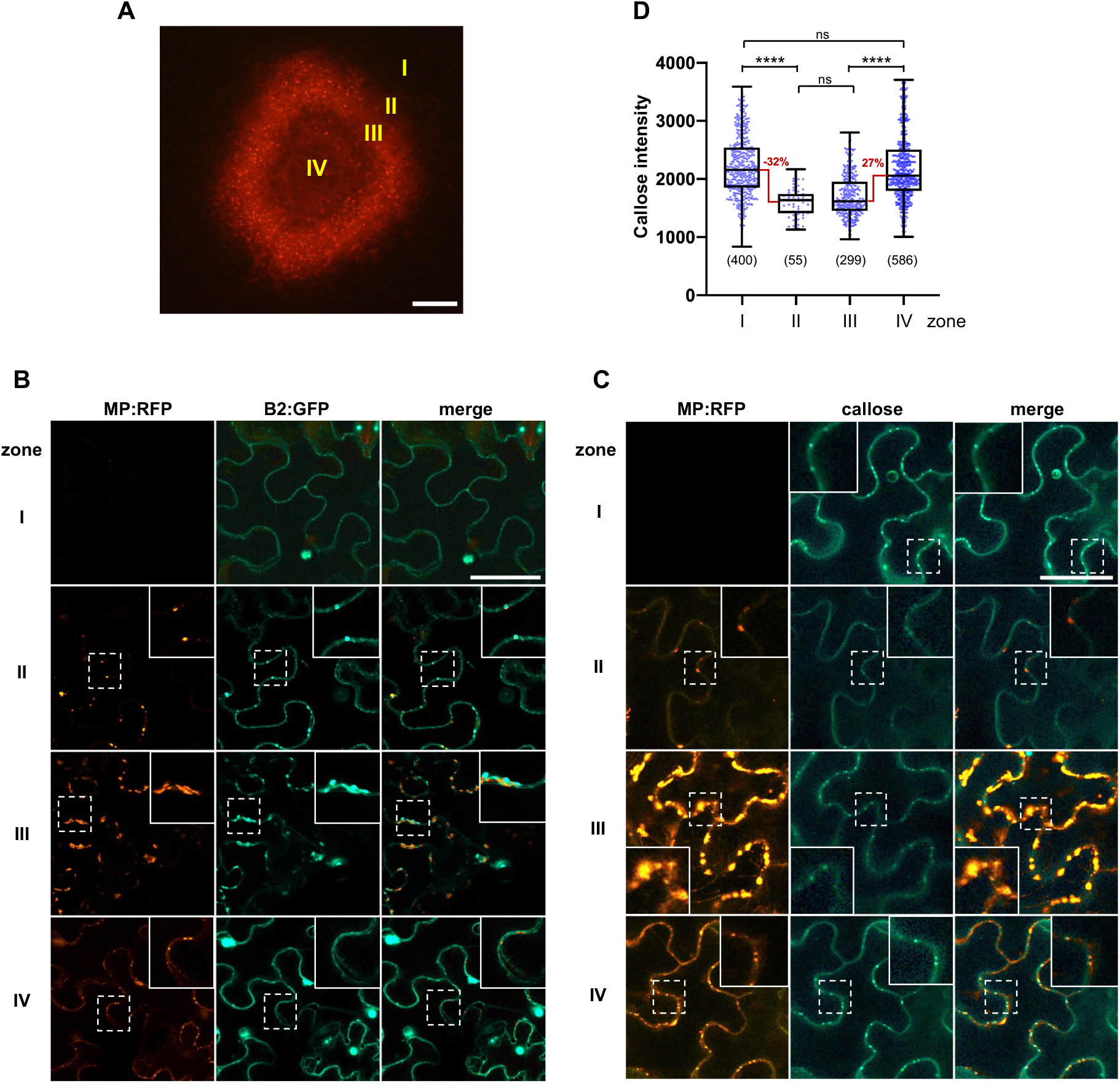
Viral MP expression correlates with a suppression of PD-associated callose levels. **(A)** Local site of infection by TMV-MP:RFP (at 4 dpi) in *N. benthamiana*. Different zones ahead of infection (zone 1), at the infection front (zone II), behind the infection front (zone III) and in the center of infection (zone IV) are indicated. Scale bar, 200 μm. **(B)** Viral dsRNA accumulation in the different zones of local TMV infection. Inlay images show magnifications of image areas framed by a dashed line. Scale bar, 20 μm. The MP of TMV is tagged with RFP (MP:RFP) and the accumulating dsRNA is imaged through binding of the *Flock house virus* B2 protein fused to GFP (B2:GFP). In cells of zone I (non-infected cells ahead of infection) B2:GFP shows a nucleo-cytoplasmic distribution, which is the typical distribution of this protein in the absence of dsRNA (Monsion et al., 2018). In cells at the virus front (zone II), B2:GFP co-localizes to MP:RFP to spots at the cell wall (likely at PD) indicating the localization of early virus-replication complexes (VRCs) engaged in virus replication and virus movement. In zone III, the VRCs have grown in size and accumulate high amounts of dsRNA consistent with high levels of virus replication to produce virus progeny. In zone IV, the MP is no longer expressed but residual MP:RFP is still seen in PD. The B2:GFP-tagged VRCs now appear rounded. (**C**) Pattern of MP:RFP and callose accumulation in the different zones. Inlays show magnifications of the image areas highlighted by dashed boxes. Scale bar, 40 μm. In zone II, where MP localizes to PD to facilitate virus movement, and to some extend also still in zone III, the PD-associated callose levels are decreased as compared to the other zones. (**D**) Quantification of PD callose in the different zones. The number of analyzed PD is shown in brackets. Two-tailed Mann-Whitney test; ****, p < 0.0001; ns, p > 0.05.

To test this hypothesis, we examined whether the expression of MP causes suppression of the poly(I:C)-induced callose deposition at PD in the absence of viral infection. Transgenic *N. benthamiana* plants that stably express MP:RFP at PD (**Figure 7, A** and **B**) complement a MP-deficient TMV mutant for movement, thus indicating that the MP:RFP in these plants is functional (**Figure 7C**). Treatment of such plants with poly(I:C) led to a 50% lower induction of callose deposition at PD as compared to WT plants (**Figure 7, D** and **E**). The ability of poly(I:C) treatment to induce callose deposition at PD was also reduced upon transient expression of MP:GFP (**Figure 7, F** and **G**). Importantly, the same effect was observed with MP^C55^:GFP. This mutant MP lacks 55 amino acids from the C-terminus but still accumulates at PD and is functional in TMV movement (Boyko et al., 2000). By contrast, dysfunctional MP^P81S^ carrying a P to S substitution at amino acid position 81, which fails to target PD and to support virus movement (Boyko et al., 2002), does not interfere with poly(I:C)-induced callose deposition. These experiments show that the TMV MP can significantly interfere with the dsRNA-induced callose deposition at PD, and that this interference requires a MP that can facilitate virus movement. Consistent with the absence of a significant role of MPK3/6 signaling in poly(I:C) induced callose deposition, expression of MP:GFP or the MP:GFP mutants did not interfere with flg22 elicitor-triggered MPK activation (**Figure 7H**). Interestingly, MP also reduces PD callose deposition induced by flg22 (**Figure 7I**), suggesting that MP interferes with signaling or signaling target mechanisms shared by both elicitors. To determine if also the MPs of other viruses interfere with the poly(I:C) induction of PD callose deposition, we tested the MPs of ORMV and turnip vein clearing virus (TVCV). The RFP-fused version of these MPs and of the MP of TMV are functional as their transient expression in *N. benthamiana* leaves allowed the intercellular spreading of the co-expressed, MP-deficient TMVΔMΔC-GFP replicon, as can be seen by the development of multiple fluorescent foci (**Figure 8A**). Consistent with function, the different MP:RFP fusion proteins colocalize with PD-associated callose (**Figure 8B**). Importantly, similar to the functional MP:RFP derived from TMV, the functional MP:RFP fusion proteins derived from ORMV and TVCV reduced the levels of PD callose deposition induced by poly(I:C) treatment (**Figure 8C**). Thus, the capacity to interfere with poly(I:C)-induced PD callose deposition may be a widespread function of viral MPs to achieve efficient infections. These observations also further substantiate the importance of antiviral PTI in the inhibition of virus movement for plants to fend off virus infections.

**Figure 7.**
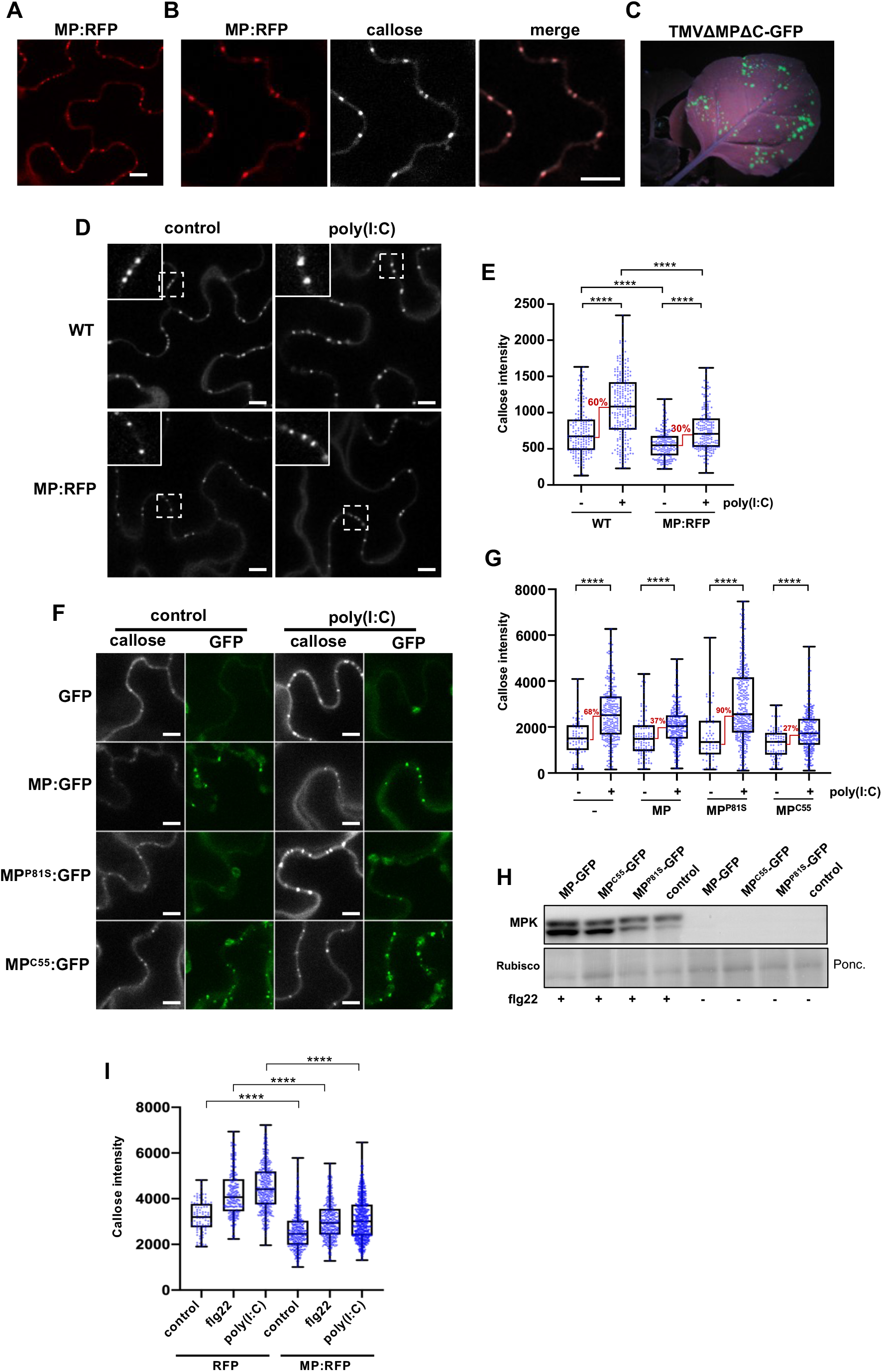
Suppression of poly(I:C)-induced immunity by MP. (**A-E**) Inhibition of dsRNA-induced callose deposition in MP:RFP-transgenic *N. benthamiana* plants. **(A**-**C**) MP:RFP is functional. (**A**) Transgenically expressed MP:RFP localizes to distinct locations at the cell wall. Scale bar, 10 μm. (**B**) The MP:RFP localizes to PD as revealed by callose staining with aniline blue. Scale bar, 10 μm. (**C**) The stably expressed MP:RFP in this line is functional and complements infection upon inoculation with *in vitro* transcribed infectious RNA of the MP-deficient TMVΔMΔC-GFP (Vogler et al., 2008), as can be seen by the occurrence of distinct GFP fluorescent infection sites at 7 dpi. (**D**) and (**E**) Inhibition of dsRNA-induced callose deposition in MP:RFP-transgenic plants. (**D**) Leaf epidermal cells of non-transgenic (WT) and MP:RFP-transgenic plants *N. benthamiana* stained with aniline blue. Inlay images show magnifications of image areas framed by a dashed line. Scale bar, 10 μm. Treatment of leaf tissues with 0.5 μg/μl poly(I:C) for 30 minutes causes a stronger increase in the level of PD-associated callose in WT plants than in MP:RFP-transgenic plants. (**E**) Quantification of callose in leaf epidermal cells upon aniline blue staining. Relative callose content in individual PD (blue dots, n > 100) as determined in three leaf discs from three plants per treatment. Two-tailed Mann-Whitney test; ****, p < 0.0001. (**F**) and (**G**) Inhibition of poly(I:C)-induced PD callose deposition by transiently expressed MP:GFP. Leaf disks excised from the GFP, MP:GFP, MP^P81S^:GFP or MP^C55^:GFP-expressing leaves 48h after agroinfiltration were incubated for one day in water, then transferred into aniline blue solution with and without 0.5 μg/μl poly(I:C) and imaged after 30 minutes. (**F**) Images of leaf epidermal cells stained for callose with aniline blue (callose) and corresponding images of the same cell area with GFP fluorescence are shown. The ability of MP:GFP to reduce the poly(I:C) induction of callose deposition at PD is inhibited by a single amino acid exchange mutation in MP (P81S) previously shown to affect its ability to efficiently target PD and to function in virus movement. Functional MP with a C-terminal deletion of 55 amino acids (C55) that targets PD also inhibits poly(I:C)-induced callose deposition like wildtype MP. (**G**) Quantification of PD-associated callose levels in leaf epidermal cells upon aniline blue staining. Leaf disks from three plants per condition were analyzed. Two-tailed Mann-Whitney test; ****, p < 0.0001. (**H**) Western blot showing that the expression of wild type or mutant MP:GFP does not interfere with flg22-induced MPK activation. Ponc., Ponceau S-stained Western blot membrane. (**I**) Inhibition of flg22- and poly(I:C)-induced PD callose deposition by MP:RFP as compared to RFP. Leaf disks excised from the RFP or MP:RFP-expressing leaves 48 hours after agroinfiltration were incubated for one day in water, then transferred into aniline blue solution with and without 0.5 μg/μl poly(I:C) and imaged after 30 minutes. Six leaf disks from two plants were evaluated for each condition. ****, p < 0.0001.

**Figure 8.**
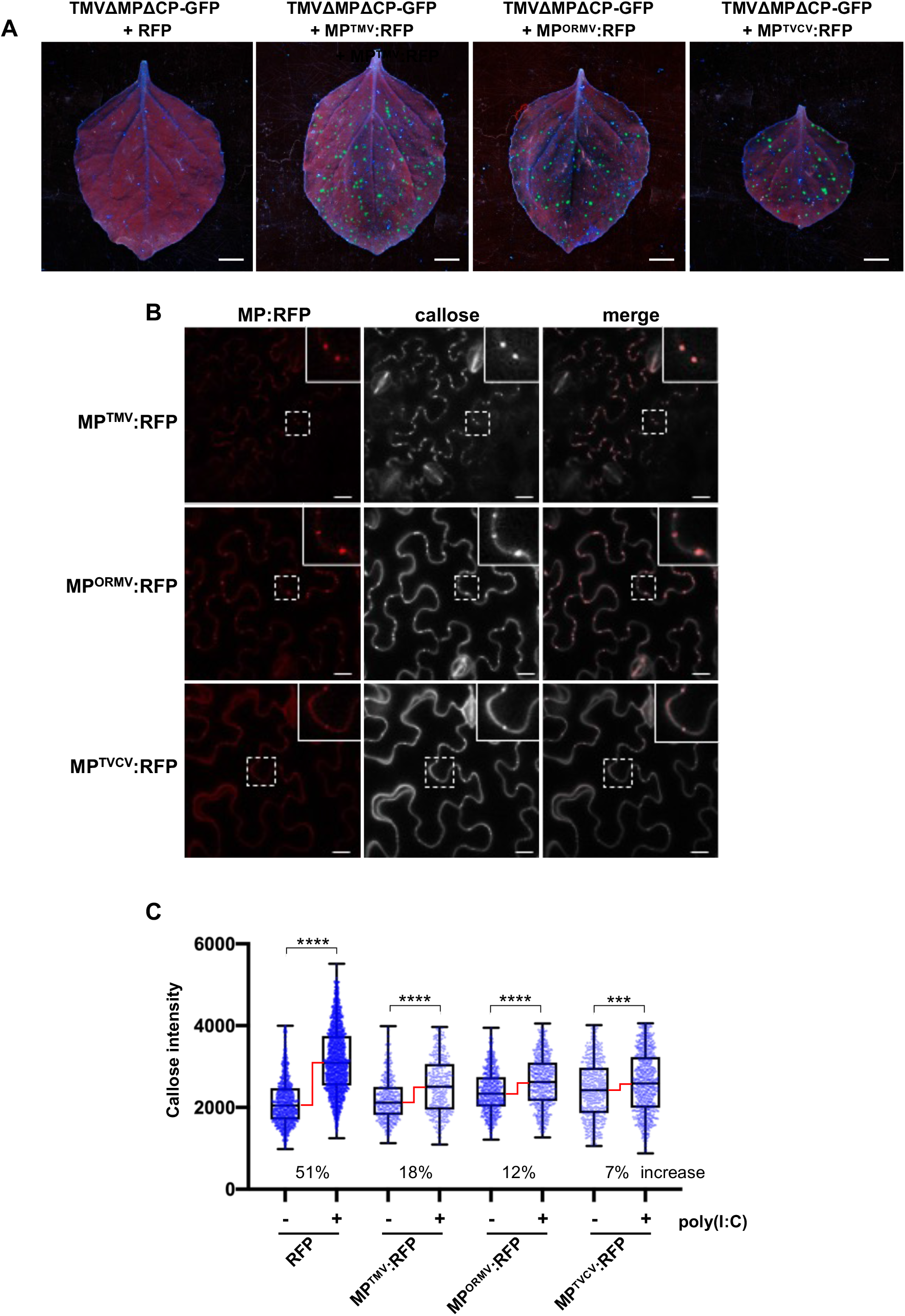
Inhibition of poly(I:C)-induced callose deposition by MPs of different viruses. (A) Different MPs are functional. Unlike free RFP, RFP fusions to the MPs of TMV (MP^TMV^:RFP),ORMV (MP^ORMV^:RFP), and TVCV (MP^TVCV^:RFP) complement the movement function of MP-deficient TMVΔM-GFP in *N. benthamiana*. Leaves were co-infiltrated with agrobacteria containing the respective RFP or MP:RFP-encoding plasmids together with highly diluted agrobacteria (OD_600 nm_ = 1 × 10^−5^) for agro-inoculation with TMVΔMPΔCP-GFP. Pictures were taken at 5 dpi. Scale bar, 1 cm. (B) MP^TMV^:RFP, MP^ORMV^:RFP, and MP^TVCV^:RFP localize to PD as shown by the presence of callose. MP-expressing leaves were stained with aniline blue and imaged after 30 minutes. Inlay images show magnifications of image areas framed by a dashed line. Scale bar, 20 μm. (C) Expression of either MP^TMV^:RFP, MP^ORMV^:RFP, or MP^TVCV^:RFP strongly reduces the induction of PD callose deposition in the presence of poly(I:C). Leaf disks excised from the RFP (control) or MP:RFP-expressing leaves 48h after agroinfiltration were incubated for one day in water, then transferred into aniline blue solution with and without 0.5 μg/μl poly(I:C) and imaged after 30 minutes. For each treatment, three images of three leaf disks taken from three plants were analyzed for PD-associated callose levels. RFP data are combined data from the 27 leaf disks that were used as RFP control in the individual agroinfiltration experiments. The increase in poly(I:C)-induced PD-callose levels seen in the presence of poly(I:C) as compared to the water-treated control is shown in percent (%). Two-tailed Mann-Whitney test; ****, p = <0.0001; ***, p = 0.0002.

### Poly(I:C) enters plant cells

As viruses replicate and produce dsRNA within cells, poly(I:C)-induced responses may only be relevant to virus infection if poly(I:C) is able to enter cells. To test this, we used B2:GFP as an intracellular dsRNA localization marker and monitored the responses of B2:GFP-transgenic *N. benthamiana* plants upon poly(I:C) treatment. Externally applied poly(I:C) may enter cells from all sides and then diffuse into the cytoplasm. Thus, a strong redistribution of B2:GFP similar as in virus-infected cells, where dsRNA production centers within the VRCs, should not be expected. Indeed, imaging GFP fluorescence under normal conditions did not show obvious changes in the distribution of B2:GFP between poly(I:C)-treated and water-treated tissues. Nevertheless, by assigning specific color to low intensity pixels, poly(I:C)-treated tissues clearly showed an accumulation of low intensity signals along the periphery of the cells (**Figure 9**), thus suggesting poly(I:C) uptake by plant cells.

**Figure 9.**
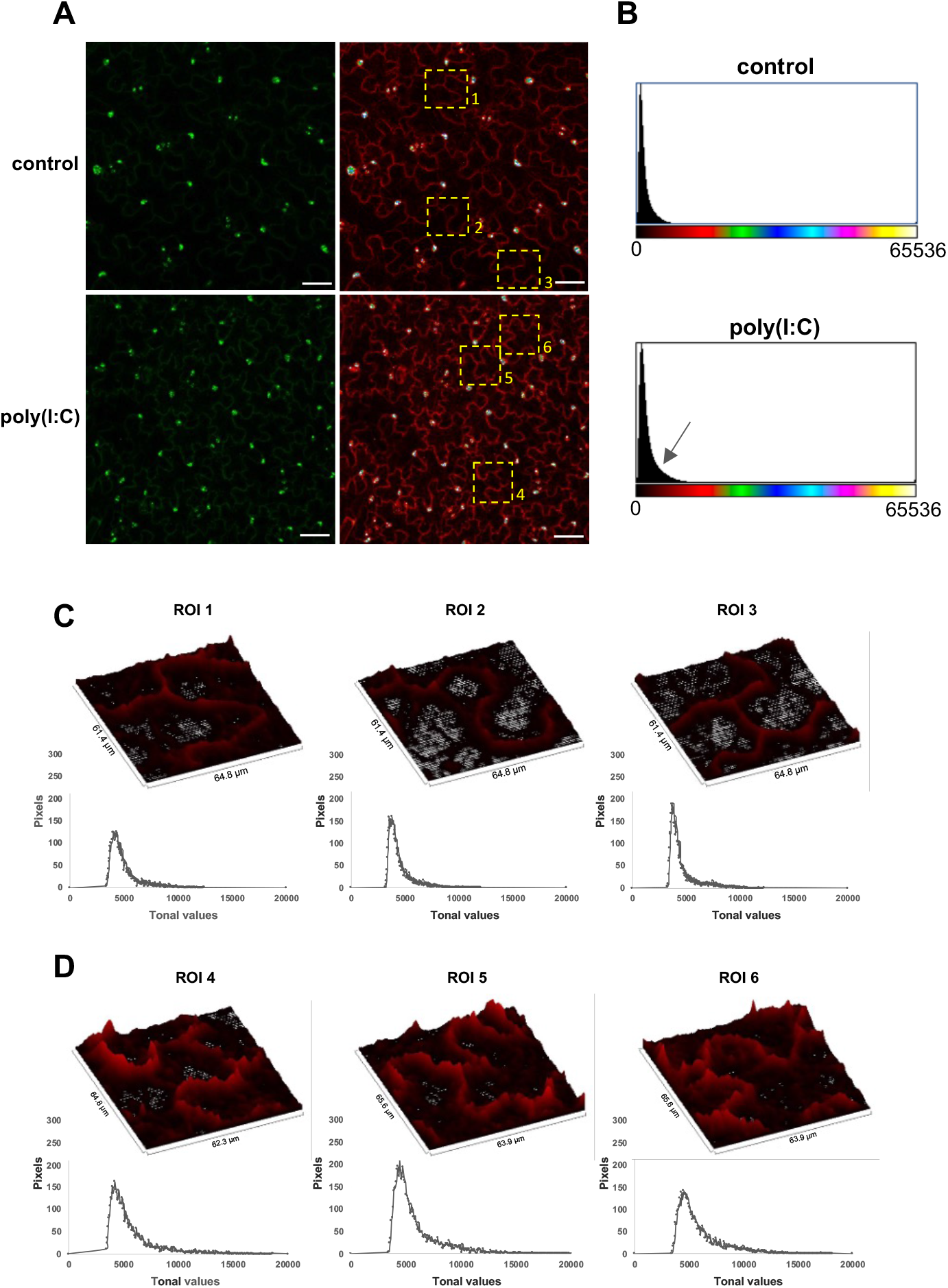
Poly(I:C) enters plant cells. **(A)** B2:GFP-transgenic *N. benthamiana* leaf tissue treated with water (control) or 0.5 μg/μl poly(I:C) and imaged with ImageJ “green” (left) and “6 shades” (right) color look-up tables (LUT). The “6 shades” LUT assigns 6 colors to specific ranges of pixel intensity values and low intensity pixels are shown in red color. As compared to the control treatment, the poly(I:C) treatment results in an enrichment of red color pixels in the periphery of the cells. Highlighted regions of interest (ROI) are further analyzed in **(C)** and **(D)**. Scale bar, 50 μm. **(B)** Histograms showing the number of pixels for each of the 65536 tonal values stored in the 16-bit “6 shades” LUT images. As compared to the control image histogram, the poly(I:C) image histogram shows an increased number of “red” pixel values. **(C)** and **(D)** Surface plot and histogram analysis of the ROIs shown in **(A)**. As compared to the surface plots of the control image ROIs, the surface plots of ROIs within the poly(I:C) image indicate a strong accumulation of B2:GFP along the periphery of the cells. This is also indicated by the corresponding histograms indicating the increased amount of red, low intensity B2:GFP pixels in the poly(I:C) ROIs as compared to the control ROIs. These observations have been confirmed by analyzing 8 images per treatment.

## Discussion

### dsRNA-induced callose deposition during virus infection

Accumulation and degradation of callose at PD play an essential role in controlling macromolecular transport between cells (De Storme and Geelen, 2014; Wu et al., 2018). Mutations and conditions that alter the levels of callose at PD strongly affect the conductivity of PD for macromolecular transport (Simpson et al., 2009; Guseman et al., 2010; Vaten et al., 2011; Benitez-Alfonso et al., 2013). A role of callose in plant-virus interactions became apparent by the observation that elevated callose levels in plants silenced for the callose-degrading enzyme restricted the spread of virus infection, thus suggesting that PD callose deposition may be part of early defense responses against virus attack (Beffa et al., 1996). However, how viruses trigger PD callose deposition and yet still maintain their cell-to-cell movement despite of this host defense response remained open. The induction of callose deposition at PD by cell-autonomous replication of an MP-deficient TMV replicon led to the conclusion that virus replication induces “stress” leading to callose deposition at PD (Guenoune-Gelbart et al., 2008), but the nature of the “stress” and the underlying mechanism remained obscure. The finding that viruses induce innate immunity (Kørner et al., 2013) and that dsRNA is a potent PAMP elicitor in plants (Niehl et al., 2016) suggests dsRNA as a potential candidate for the perceived stress signal. Our data shown here that dsRNA-induced immunity is linked to PD callose deposition raise a model that virus replication causes callose deposition and PD closure mediated through a PTI response triggered by viral dsRNA.

### MP facilitates virus movement by suppressing a dsRNA-induced PTI response

Because virus movement depends on prior replication of the viral genome (Christensen et al., 2009), and given that the virus must continue to replicate to produce progeny, the PTI response is likely triggered immediately in the newly infected cells at the infection front and maintained in cells behind the front. Thus, the perception of dsRNA may target PD for closure throughout the infection site. This mechanism may have evolved to isolate the infected cells from surrounding cells to prevent further spread of infection but also to protect the virus replication and virus progeny production in the infected cells against intercellular defense signaling. At the infection front, the dsRNA-producing virus must nevertheless be able to overcome PD closure in order to spread infection into non-infected cells. Consistently, pioneering studies with TMV showed that the virus moves between cells with the help of virus-encoded MP, that MP targets PD, and that it increases the size exclusion limit (SEL) of PD and thus the permeability of the channels for macromolecular trafficking (Citovsky, 1999). Importantly, using microinjection, MP was shown to gate the PD between cells only at, but not behind, the virus infection front (Oparka et al., 1997). Our observations link this activity to the suppression of a dsRNA-induced response by showing (i) that cells at the virus infection front, where MP increases the PD SEL, have a significantly lower level of callose at PD as compared to other cells and (ii) that the induction of PD callose deposition by dsRNA [poly(I:C)] is significantly reduced by ectopically expressed MP. The ability of MP of TMV to interfere with dsRNA-induced callose deposition seems to reflect an activity shared with other MPs as similar to MP^TMV^ also the expression of MP^ORMV^ or MP^TVCV^ reduced the intensity of PD callose induction by poly(I:C) as compared to the control (absence of the respective MP) (**Figure 8**). These observations suggest a new paradigm for virus movement whereby dsRNA produced by TMV replication in a newly infected cell at the infection front triggers a host PTI response that targets PD for callose deposition and closure in order to restrict the spreading of the virus. To allow the spread of replicated viral genomes into non-infected cells, the viral MP acts as an effector to transiently suppress this dsRNA-induced response (**Figure 10, A** and **B**). Upon enhancement of the PTI response by external treatment of infected plants with poly(I:C) the virus-encoded MP may become insufficient for efficient suppression, thus leading to the inhibition of virus movement (**Figure 1, A** and **B**).

**Figure 10.**
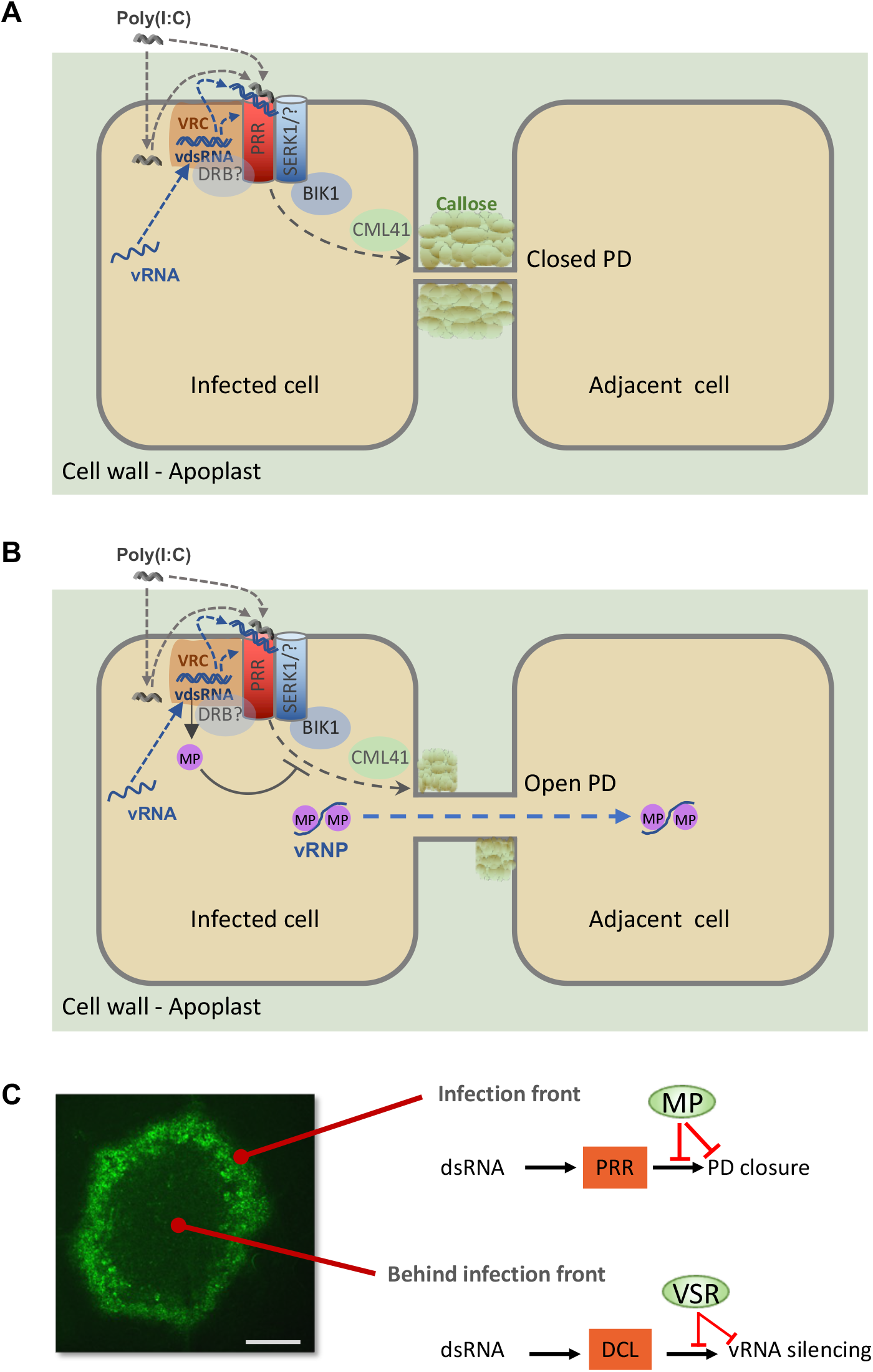
Virus infection facilitated by virus-encoded effector proteins. (**A**) and (**B**) Suppression of PTI by MP. (**A**) Perception of dsRNA produced in cortical ER-associated VRCs at the PM by an unknown cytoplasmic or membrane-associated pathogen recognition receptor (PRR) and the SERK1 co-receptor (with potential contribution by one or more other co-receptors) triggers a signaling pathway leading to callose deposition and PD closure. dsRNA produced during infection may require secretion into the apoplasm to allow perception at the PM (dashed blue lines and arrows). Externally applied poly(I:C) may be perceived from the apoplasm or secreted upon initial uptake by the cells (dashed gray lines and arrows). (**B**) MP suppresses dsRNA-triggered callose deposition and allows intercellular spread of the viral ribonucleoprotein complex (vRNP). The MP may interact with intracellular PTI signaling as indicated or interact with callose synthesizing or degrading enzymes at PD. The MP may also be secreted to inhibit dsRNA perception at the PM. (**C**) dsRNA triggers PTI and antiviral RNA silencing and both responses are suppressed by viral effector proteins to support virus propagation. Whereas MP acts in cells at the virus front to facilitate virus movement by blocking a dsRNA-induced callose defense response at PD, the VSR blocks dsRNA-induced antiviral RNA silencing in the center of infection sites to support virus replication and production of virus progeny. A local infection site of TMV encoding MP fused to GFP (TMV-MP:GFP, 7 dpi) in *N. benthamiana* is shown. Scale bar, 1 mm.

The MP is expressed in cells at and also closely behind the infection front (**Figure 6, A** to **C**). The restriction of MP activity and of low callose levels in cells at the infection front indicates that the protein exists in different activity states. Consistently, several studies correlated the activity of MP with its phosphorylation state (Lee and Lucas, 2001). The partial, rather than full, suppression of dsRNA-induced callose deposition by ectopically expressed MP may reflect the different activity states of MP expressed under such conditions.

The mechanism by which MP suppresses the dsRNA-induced callose deposition at PD remains to be studied. dsRNA sequestration by MP is precluded as MP has no dsRNA binding activity (Citovsky et al., 1990). Electron micrographs indicated that the TMV MP forms fibrillar substructures within PD cavities (Ding et al., 1992; Moore et al., 1992; Heinlein et al., 1998) but it is unknown whether these structures are functional. Several studies support the hypothesis that a callose-degrading beta-1,3-glucanase enzyme activity may regulate virus movement (Iglesias and Meins, 2000; Bucher et al., 2001; Levy et al., 2007), but strong evidence indicating that viruses like TMV indeed operate such activities for virus movement is lacking. More recent observations suggest that the MPs of different viruses interact with the synaptotagmin SYTA for movement (Uchiyama et al., 2014; Levy et al., 2015; Yuan et al., 2018). Synaptotagmins (SYTs) and other Ca^2+^-sensitive C2 domain-containing proteins, such as the multiple C2 domains and transmembrane region proteins (MCTPs) are proposed to function as membrane-tethering proteins at membrane contact sites (Tilsner et al., 2016; Brault et al., 2019). Notably, the strand of endoplasmic reticulum (desmotubule) that traverses the PD channel between cells is tethered to the adjacent plasma membrane (PM) by MCTPs and SYTs (Brault et al., 2019; Ishikawa et al., 2020). Therefore, it is conceivable that MPs target SYTA to reach PD or even to modify a membrane tethering activity of SYTA within PD in order to alter the cytoplasmic space between the tethered membranes available for macromolecular transport (Pitzalis and Heinlein, 2017). However, while further studies are needed to explore this idea, the results shown here promote the model that the MPs of TMV and also the MPs of ORMV and TVCV facilitate movement by interacting with components of the dsRNA-induced signaling and callose synthesis and turnover pathways to inhibit callose deposition at PD. The MPs may interfere with these pathways in the cytoplasm (**Figure 10A and B**) or at PD. However, earlier studies indicated that the MPs of TMV, TVCV and cauliflower mosaic virus have the capacity to interact with the cell wall protein pectin-methylesterase (PME) which may allow transport of MP through the secretory pathway (Chen et al., 2000). It is conceivable, therefore, that MPs are partially secreted to inhibit the activity of the dsRNA perceiving receptors at the PM. However, whether the MPs of TMV, ORMV and TVCV inhibit PD callose deposition through interaction with PTI signaling components or rather through direct interactions with callose synthesizing or degrading enzymes at PD remains to be investigated. As the MP of TMV suppressed PD callose deposition triggered by either poly(I:C) and flg22, the signaling components, enzymes, or mechanisms affected by MP likely play a role in both poly(I:C)- and flg22-triggered pathways.

### dsRNA induces antiviral defense through a novel PTI signaling pathway

We have shown here that the dsRNA-induced signaling pathway leading to callose deposition at PD involves SERK1, BIK1/PBL1, CML41, Ca^2+^, and potentially PDLP1/2/3, but neither strong MPK activation or ROS production nor the presence of the major ROS-producing enzymes (RBOHD and RBOHF). Thus, although MPK activation and ROS production are hallmarks of PTI (DeFalco and Zipfel, 2021) and ROS signaling plays a role in PD regulation (Cheval and Faulkner, 2018), dsRNA activates its own specific pathway to regulate PD. Moreover, the lack of ROS production clearly distinguishes virus/dsRNA-induced signaling from the ROS-associated responses induced by other pathogens. Importantly, we found that poly(I:C) treatment triggers BIK1 phosphorylation and that the level of poly(I:C)-induced BIK1 phosphorylation depends on SERK1, thus potentially suggesting the existence of dsRNA-perceiving SERK1-containing receptor complex that signals to BIK1. Alternatively, dsRNA may induce yet non-identified signaling molecules that are signaled through SERK1 and BIK1. The PD-localized CML41 protein was previously shown to participate in flg22-triggered, but not chitin-triggered PD callose deposition (Xu et al., 2017). Thus, although differing in upstream components, dsRNA-induced signaling may target PD-associated regulatory components also regulated by flg22. The absence of ROS signaling in dsRNA-induced PD regulation could reflect this specific elicitor type or a specific location of perception. Bacterial and fungal PAMPs are released in the apoplast and perceived by PRRs at the PM, while viral dsRNA formation and perception may occur at intracellular membranes where viruses replicate. RIG-I-like receptors (RLRs) including RIG-I, MDA5 and LGP2 detect the presence of dsRNA in animals (Tan et al., 2018) and provide examples of dsRNA perception in the cytoplasm. Viral RNAs in animals are also recognized by Toll-like receptors in endosomes following internalization by dendritic cells and macrophages. In analogy, it is known that bacterial PAMP receptor complexes in plants are internalized from the PM to endosomes and that plant receptor-complexes can signal from endosomes. Recognition of viral dsRNA may therefore occur during membrane fusion events between viral RNA-containing vesicles or membrane-associated viral replication complexes (VRCs) (Niehl and Heinlein, 2019). Alternatively, the cytoplasmic dsRNA may be exported into the apoplast to be sensed by the PM-resident PRRs. A precedent for the transport of functional RNA molecules across plant membranes is provided by cross-kingdom RNAi (Huang et al., 2019). For example, dsRNAs sprayed onto plants were shown to inhibit fungal growth in tissues distant from the sprayed tissues by inducing RNAi against essential fungal genes thus suggesting that applied dsRNAs are taken up by plants and that either dsRNAs or derived small RNAs processed within cells are able to reach the fungus in distant tissues (Koch et al., 2016). Moreover, emerging evidence suggest the presence of viral particles, viral proteins, and viral RNA in the apoplast of infected plants (Mohaved et al., 2019; Hu et al., 2021; Wan et al, 2015; Wan and Laliberté, 2015). Moreover, extracellular vesicles that are secreted from plants cells and are present in apoplastic fluid contain various RNA species, including viral RNA (Cai et al., 2018; Cai et al., 2021; Ruf et al., 2022). Poly (I:C) used in this study mimics these potential pathways by apparently being able to enter plant cells upon treatment (**Figure 9**), which is consistent with dsRNA perception in the cytoplasm, or be sensed in the apoplasm, either upon secretion or before cell entry (**Figure 10A** and **B**). Plant viruses like TMV and ORMV undergo early replication stages at punctate, cortical microtubule-associated ER sites in close vicinity of the PM (potentially ER:PM contact sites (Pitzalis and Heinlein, 2017; Huang and Heinlein, 2022)), which may facilitate dsRNA perception through membrane fusion events or dsRNA secretion from the VRCs and activation of PM-localized signaling proteins in the apoplasm (Niehl and Heinlein, 2019). DRB2 and other double-stranded RNA binding proteins (DRBs) were recently shown to accumulate in VRCs and to play a role in virus accumulation (Incarbone et al., 2021) or virus-induced necrosis (Fatyol et al., 2020) and could have an important function in dsRNA sensing. Importantly, dsRNA-induced innate immunity is unaffected by mutations in dsRNA binding DICER-LIKE (DCL) proteins, which excludes these proteins as the dsRNA receptors for PTI and also shows that dsRNA silencing and dsRNA-induced innate immunity require different protein machinery (Niehl et al., 2016). The two different antiviral defense responses are also spatially separated. Inactivating the TMV VSR causes virus silencing but has no effect on virus movement and causes potent antiviral silencing only in cells behind the front engaged in virus replication for the production of virus progeny (Kubota et al., 2003; Vogler et al., 2007). Thus, we propose a model whereby a virus requires MP as viral virulence effector in cells at the infection front to suppress dsRNA-triggered PTI to support virus movement whereas a VSR is required as virulence effector in cells behind the front to suppress dsRNA-induced silencing for producing viral progeny (**Figure 10C**).

It will be interesting to dissect how the SERK1-BIK1/PBL1 module signals to CML41 and PDLP1/2/3 to regulate PD callose deposition. It is possible that BIK1/PBL1 may interact with and phosphorylate directly PDLPs in mediating callose deposition at the PD. It has been shown that the plasma membrane-tethered BIK1 regulates PAMP-triggered calcium signals by directly phosphorylating cyclic nucleotide-gated channel 2 and 4 (CNGC2/4), whose activities can be regulated by CAM7 (Tian et al., 2019). Additionally, BIK1 also phosphorylates calcium-permeable channel, hyperOsmolality-induced [Ca^2+^] increase 1.3 (OSCA1.3) in regulating stomatal immunity (Thor el al., 2020). Our observation that the calcium-chelating EGTA inhibits dsRNA-induced PD callose deposition supports the involvement of the calcium binding protein CML41 and calcium signaling in PD regulation. Therefore, it will be interesting to determine if any calcium channels directly regulated by BIK1/PBL1 and modulated by CML41 may be involved in mediating calcium signaling in plant antiviral immunity. This will help to delineate a genetic and biochemical signaling pathway linking SERKs-RLCKs-calcium channels/CML41-PDLP1/2/3 to calcium signals in regulating PD and plant antiviral immunity. It will also be interesting to determine the role of other SERKs in virus sensing and immunity. It has been shown that SERK3/BAK1 plays a role in antiviral defense (Kørner et al., 2013) but the molecular mechanism remains to be explored. While poly(I:C)-induced ethylene production was not affected in *serk3/bak1* mutants (Niehl et al., 2016), it remains to be tested if SERK3/BAK1 or also SERK2 or SERK4 could play a role in dsRNA sensing leading to PD callose deposition, thus potentially explaining cases of *serk1*-independent dsRNA-induced gene activation (**Figure 2A**) as well as the residual poly(I:C)-induced activation of PD callose deposition in *serk1* mutants (**Figure 3E**).

In conclusion, we found that as a plant defense mechanism, dsRNA-induced antiviral PTI targets PD for callose deposition through some shared typical PTI signaling components with distinct features. To counteract this and launch efficient infections, viral MPs could effectively suppress dsRNA-induced callose deposition at PD, thus leading to a new concept of plant-virus interaction arm-race. This study calls upon the identification of the PTI dsRNA receptor, the mechanisms of SERK1-BIK1-calcium channels/CML41-PDLP1/2/3 signaling and PTI suppression by MP, and how dsRNA-induced PTI and RNA silencing are controlled during the spread of infection, all of which present exciting new challenges for additional studies.

## Methods

### Plant materials and growth conditions

*N. benthamiana* and *A. thaliana* plants were grown from seeds in soil. *N. benthamiana* were kept under 16h/8h light/dark periods at +22 °C/+18 °C in a greenhouse equipped with Philips SON-T 400 W HPS Lamps (200-250 μmol/m^2^/s). *A. thaliana* plants were kept under 12h/12h light/dark periods at +21°C/+18 °C in a growth chamber equipped with LED lights (160-175 μmol/m^2^/s). MP:RFP-transgenic *N. benthamiana* plants were produced by leaf disk transformation (Horsch et al., 1985) using binary plasmid pK7-MP:RFP (Boutant et al., 2010). The plasmid was constructed by inserting the MP coding sequence of TMV into pK7RWG2 using Gateway procedures. *N. benthamiana* plants expressing GFP fused *Flock house virus* B2 protein have been described previously (Monsion et al., 2018) and were provided by Christophe Ritzenthaler (IBMP, CNRS, Strasbourg, France). The Arabidopsis mutants used in this study have been described previously and homozygous seeds were kind provided from different research laboratories. Seeds of the *bik1 pbl1* double mutant (SALK_005291 SAIL_1236_D07) (Zhang et al., 2010) were provided by Cyril Zipfel (University of Zürich, Switzerland). *mpk3-1* (SALK_151594) and *mpk6-2* (SALK_073907) lines were given by Kenichi Tsuda (Max Planck Institute for Plant Breeding research, Cologne, Germany) and, together with seeds of the *mpk3amiRmpk6* (Li et al., 2014) and *serk1-1* (SALK_044330) (Meng et al., 2015b) mutants, also by the author Libo Shan. Seeds for *rbohd* and *rbohf* mutants (Torres et al., 2002) were provided by Christine Faulkner (John Innes Centre, Norwich) and seeds of *CML41* overexpressing and silenced lines (*CML41-OEX-2, CML41-OEX-12, CML41-amiRNA-1* and *CML41-amiRNA-4*) (Xu et al., 2017) were a gift of Matthew Gilliham (University of Adelaide, Australia). Arabidopsis lines transgenic for PD markers mCherry-BDCB1 (Simpson et al., 2009; Benitez-Alfonso et al., 2013) or PdBG2-citrine (Benitez-Alfonso et al., 2013) were provided by Yoselin Benitez-Alfonso (University of Leeds, UK).

### Virus inoculation

cDNA constructs for TMV-MP:RFP (Ashby et al., 2006), TMV-GFP (Heinlein et al., 1995), and TMVΔM-GFP (Vogler et al., 2008) have been described previously. *N. benthamiana* plants were mechanically inoculated in the presence of an abrasive (Celite®545) with infectious RNA *in vitro*-transcribed from these constructs. A TMV replicon (TMV-ΔMP-ΔCP-GFP) cloned in a binary vector for agroinfiltration and used for testing virus replication and movement trans-complementation has been described (Borniego et al., 2016). Arabidopsis plants were inoculated with purified ORMV virions (Niehl et al., 2012).

### Analysis of virus infection in the presence of elicitors

To test the effect of elicitors on TMV-GFP infection in *N. benthamiana*, plants were inoculated with 200 μl inoculum containing 20 μl of infectious viral RNA transcription mix and 0.5 μg/μl (equals ~1 μM) poly(I:C) (Sigma-Aldrich, USA), or 1 uM flg22 (EZBiolabs, USA or Proteogenix, France), or water. Infection sites on the inoculated leaves were imaged using a hand-held camera and UV lamp (BLAK RAY B-100AP; UVP Inc., Upland, California) in the presence of a ruler for size normalization. The areas of infection sites in each leaf were measured with Image J software upon selection of infection site as regions of interest using fluorescence thresholding and the wand tracing tool, and by setting the scale according to the ruler. For testing the effect of poly(I:C) on viral replication, *N. benthamiana* leaves were agro-inoculated with a MP-deficient, cell-autonomous TMV replicon (TMVΔM-ΔC-GFP) (Borniego et al., 2016). After 1 day, the fluorescent leaf patches were gently rubbed with 200 μl of 0.5 μg/μl poly(I:C) in the presence of celite. At 1, 3 and 5 days after this treatment, the GFP-expressing leaf patches were analysed for viral RNA accumulation by quantitative Taqman RT-qPCR using previously described methods (Mansilla et al., 2009; Niehl et al., 2016).

To test the effect of elicitors on ORMV infection in Arabidopsis, 4 μl of elicitor solution (10 μg/ml poly(I:C) or 10 μM flg22) or 4 μl PBS were placed on rosette leaves of 3 weeks old Arabidopsis wildtype or mutant Col-0 plants. A volume of 2.5 μl of a 20 ng/μl solution of purified ORMV virions was placed on the same leaves. Subsequently, the leaves were gently rubbed in the presence of celite as abrasive. Immediately after treatment, remaining elicitors, buffers and virions were washed off the leaf surface. Symptoms were analysed at 28 dpi. At the same time young, systemic leaves were sampled for analysis of virus accumulation by quantitative Taqman RT-qPCR using previously described methods (Mansilla et al., 2009; Niehl et al., 2016).

### Analysis of differential gene expression by RT-qPCR

*N. Benthamiana* or Arabidopsis Col-0 leaf discs were excised with a cork borer and incubated overnight in 12-well plates containing 600 μl deionized, ultra-pure water. The leaf disks were washed several times with water and then incubated with elicitor (1 μM flg22, 0.5 μg/μl poly(I:C), or water as control) for 3 hours. After washing the discs with deionized, ultra-pure water three times, samples were ground to a fine powder in liquid nitrogen and total RNA was extracted by TRIzol™ reagent according to the protocol of the manufacturer. 2 μg of RNA were reverse transcribed using a reverse transcription kit (GoScript™ Reverse Transcription System, Promega). The abundance of specific transcript was measured by probing 1 μL cDNA by quantitative real-time PCR in a total volume of 10 μl containg 5 μL SYBR-green master mix (Roche), 0.5 μM forward and reverse primer and water. qPCR was performed in a Lightcycler480 (Roche) using a temperature regime consisting of 5 minutes at 95°C followed by 45 cycles at 95°C for 10 seconds, 60°C for 15 seconds, and 72°C for 15 seconds, and ending with a cycle of 95°C for 5 seconds, 55°C for 60 seconds, 95°C for continuous time until final cooling to 40°C for 30 seconds. The threshold cycle (CT) values were normalized to CT-values obtained for reference genes ACTIN 2 and UBIQUITIN 10 (Czechowski et al., 2005), providing ΔCT values. These were used to calculate the 2^−ΔCT^ values representing relative expression levels, the mean values and standard errors (SE). Each mean value represents the analysis of three independent replicate samples (individual plants treated the same way and harvested at the same time), each measured by three technical replicates. Primers are listed in Table S1.

### Transient expression of proteins by agroinfiltration

Binary plasmids for transient expression of RFP and of TMV-derived MP:RFP, MP:GFP, MP^C55^:GFP, and MP^P81S^:GFP as well as of MP^ORMV^:RFP and MP^TVCV^:RFP were created by Gateway cloning as has been described previously (Brandner et al., 2008; Sambade et al., 2008; Boutant et al., 2010).

For transient expression of the fluorescent fusion proteins, cultures of *A. tumefaciens* bacteria (strain GV3101) carrying these plasmids were harvested by centrifugation, resuspended in infiltration medium (10 mM MES, 10 mM MgCl_2_, 200 μM acetosyringone; pH 5.5) to a final optical density at 600 nm (OD_600_) of 0.1 (unless stated differently), and infiltrated into the abaxial side of the leaf using a syringe without a needle. Leaves were observed by confocal microscopy at 48 hours after agroinfiltration.

For GFP mobility assays, we used Agrobacteria that were co-transformed with binary vectors for expression of GFP together with the cell-autonomous nuclear protein NLS:RFP (pB7-NLS:M2CP:RFP; this vector was created by recombining pZeo-NLS:MS2CP (Sambade et al., 2008) with expression vector pH7RWG2). The two binary vectors carry different resistance genes and their presence in the same agrobacteria was maintained by appropriate antibiotic co-selection. Before infiltration, the diluted culture (OD_600_ = 0.1) was further diluted 1:1000 or 1:10000 to ensure expression of both proteins in only few cells of the leaf. 24 hours after infiltration, the agroinfiltrated leaves were detached and analyzed by confocal microscopy, revealing about 10-15 single cell transformation events with both markers. The levels of cytoplasmic GFP and nuclear NLS:RFP differed between the transformed cells but cells expressing only GFP or only NLS:GFP were not observed, which excludes the occurrence of transformation events in which only one of the T-DNAs was transferred. Subsequently, leaf disks with the single cell transformation events were excised and incubated in 0.5 μg/μl poly(I:C) or water for 48 hours. Finally, each of the transformation events was evaluated for GFP movement by confocal microscopy by counting the radial cell layers into which GFP has moved away from the infiltrated cell (marked by red fluorescent nucleus).

For movement trans-complementation assays, *N. benthamiana* leaves were infiltrated with *Agrobacterium* cultures (OD_600 nm_ = 0.3) for the expression of either MP^TMV^:mRFP MP^ORMV^:mRFP, MP^TVCV^:mRFP or of free mRFP together with a highly diluted *Agrobacterium* culture for infection of single cells with TMVΔMPΔCP-GFP (OD_600 nm_ = 1 × 10^−5^). Fluorescent infection sites indicating complementation of the MP-deficient virus were imaged with a Nikon D80 camera at 5 dpi under UV illumination. For the movement trans-complementation assay with MP:RFP-transgenic *N. benthamiana* plants, leaves were inoculated with infectious RNA *in vitro*-transcribed from pTMVΔM-GFP (Vogler et al., 2008) and infection sites were observed at 7 dpi.

### Callose staining

Leaf disks were excised with a cork borer and placed into wells of 12-well culture plates containing 1 ml water and incubated overnight under conditions at which the plants were raised. The leaf discs were washed several times with water before use. For callose staining, individual leaf disks were placed on microscope slides and covered with a coverslip fixed with tape. If not otherwise stated, 200 μl of a 1% aniline blue solution (in 50 mM potassium phosphate buffer, pH 8.0) containing either 0.5 μg/μl poly(I:C), 50 ng/μl dsRNA^phi6^ or 1 μm flg22 were soaked into the space between the glass slide and coverslip. The glass slide with the sample was evacuated for 1-2 minutes (< 0.8 pa) in a vacuum desiccator followed by slow release of the pressure. Aniline blue fluorescence was imaged 30 minutes after dsRNA or control treatment using a Zeiss LSM 780 confocal laser scanning microscope with ZEN 2.3 software (Carl Zeiss, Jean, Germany) and using a 405 nm diode laser for excitation and filtering the emission at 475-525 nm. 8-bit Images acquired with a 40× 1.3 N.A. Plan Neofluar objective with oil immersion were analyzed with ImageJ software (http://rsbweb.nih.gov/ij/) using the plug-in *calloseQuant*, which after setting few parameters localizes fluorescent callose spots and quantifies callose fluorescence intensity of each spot automatically (Huang et al., 2022). This plugin is available at https://raw.githubusercontent.com/mutterer/callose/main/calloseQuant_.ijm. Callose spots were measured in 1-3 images taken from each leaf disk. If not otherwise mentioned, three leaf discs from three different plants were evaluated for each genotype or condition. To control for normal poly(I:C) treatment and callose staining conditions, samples of Arabidopsis mutants were always analyzed in parallel to samples from the Col-0 wild-type. Similarly, samples from agroinfiltrated MP:GFP/RFP-expressing *N. benthamiana* leaves were compared with samples from agroinfiltrated GFP/RFP-expressing control leaves. The distribution of pooled fluorescence intensities obtained for the specific genotype or treatment condition is shown in boxplots or column diagrams. Regions of interest (ROIs) selected by *calloseQuant* were verified visually before measurement. ROIs that were not at the plant cell walls or showed no clear signal above the background signal were deleted. Moreover, individual fluorescence intensities that occurred as outliers from the general distribution of fluorescence intensities (<1%) in the sample were excluded from analysis.

### Analysis of MPK activation

Leaf disks from 4-week-old *A. thaliana* or *N. benthamiana* plants were elicited with 1 μM flg22 or 0.5 μg/μl (equals ~1 μM) poly(I:C). As a control, leaf discs were treated with water or treated with PBS. Elicitor and control treatment was performed by addition of the elicitor or the controls to leaf disks acclimated overnight in ultrapure water. After addition of the elicitor, leaf disks were vacuum infiltrated for 10 min. Samples were taken after an additional 20 min of incubation. MPK phosphorylation was determined using protein extracts obtained from elicitor or control-treated leaf disks using immunoblots probed with antibodies against phosphor-p44/42 ERK (Cell Signaling Technology, Beverly, MA, USA; Ozyme S4370S) and horseradish peroxidase (HRP)-labelled secondary antibodies for luminescence detection (SuperSignal™ West Femto Maximum Sensitivity Substrate, ThermoFisher, France).

### Analysis of ROS production

Leaf discs excised from 4-week-old *A. thaliana* or *N. benthamiana* plants were incubated overnight in 96-well plates with 600 μL of deionized, ultra-pure water. The next day deionized, ultra-pure water was replaced with 100 μL reaction solution containing 50 μM luminol and 10 μg/mL horseradish peroxidase (Sigma, USA) together with or without 1 μM flg22 or 0.5μg/μl poly(I:C). Luminescence was determined with a luminometer (BMG LABTECH, FLUOstar®Omega) at 1.5 minute intervals for a period of 40 minutes. Mean values obtained for 10 leaf discs per treatment were expressed as mean relative light units (RLU).

### Seedling growth inhibition assay

Seeds were surface-sterilized and grown vertically at 22°C under 12h/12h light/dark periods in square petri-dishes on half-strength Murashige and Skoog (MS) basal medium (pH 5.8) containing 0.5 g/L MES and 0.8% agar. 7 days old seedlings were transferred into liquid half-strength Murashige and Skoog (MS) medium with or without 500 ng/μl (equals ~1 μM) poly(I:C) or 1 μM flg22. The effect of treatment on seedling growth was documented on photographs 12 days after treatment and measured with a ruler.

### Protoplast transient expression and BIK1 mobility shift assays

Arabidopsis protoplasts (about 40,000 cells) isolated from wildtype Col-0 or *serk1-1* were transfected with HA-epitope-tagged BIK1 (pHBT-35S::BIK1-HA) or co-transfected with pHBT-35S::BIK1-HA and pHBT-35S::SERK1-FLAG. Protoplast isolation and the transient expression assay were done as described previously (He et al., 2007). Also the BIK1 and SERK1 constructs have already been described (Lu et al., 2010; Meng et al., 2015a). The transfected protoplasts were incubated at room temperature overnight. After stimulation with flg22 (1 μM) or poly(I:C) (0.5 μg/ul) for 20 minutes, the protoplasts were collected by centrifugation and lysed by vortexing in 100 μl co-IP buffer (150 mM NaCl, 50 mM Tris-HCl, pH7.5, 5 mM EDTA, 0.5% Triton, 1 × protease inhibitor cocktail. Before use, 2.5 μl 0.4 M DTT, 2 μl 1 M NaF and 2 μl 1 M Na_3_VO_3_ were added per 1 ml IP buffer). A final concentration of 1 μM K-252a inhibitor (Sigma-Aldrich, 05288) was added 1 hour before poly(I:C) (0.5 μg/ul) treatment. Lysed protoplasts were treated with calf intestinal phosphatase (CIP) (New England Biolabs) for 60 minutes at 37°C (1 unit per μg of total protein). BIK1 was detected in Western blots assays using HA-HRP antibody (Invitrogen, 26183-HRP)) and its phosphorylation was quantified by calculating the ratio between the intensity of the shifted upper band of phosphorylated BIK1 (pBIK1) and the sum of the intensities of both shifted and non-shifted bands (pBIK1 + BIK1) (no treatment set to 0.0). The BIK1 mobility shift assays have been described previously (Lu et al., 2010; Ma et al., 2020). SERK1-FLAG was detected in Western blot assays using monoclonal anti-FLAG (Sigma-Aldrich, F1804).

### Imaging

Microscopical imaging was performed with a Zeiss LSM 780 confocal laser scanning microscope equipped with ZEN 2.3 software (Carl Zeiss, Jean, Germany). Excitation / emission wavelengths were 405 nm/475-525 nm for aniline blue, 488 nm / 500-525 nm for GFP, and 561 nm/560-610 nm for RFP.

## Acknowledgements

We acknowledge funding from the Agence National de la Recherche (Plant-KBBE2012 / ANR-13-KBBE-0005-01, ERA-NET SusCrop2 / ANR-21-SUSC-0003-01; ANR-PRC / ANR-21-CE20-0020-01) and the Chinese Scholarship Council (PhD fellowship for CH) to MH, and funding from the NIH (R35GM144275), NSF (IOS-2049642), and the Robert A. Welch Foundation (A-2122-20220331) to LS. We thank Cyril Zipfel, Christine Faulkner, Matthew Gilliham, Kenichi Tsuda, Christophe Ritzenthaler, and Yoselin Benitez-Alfonso for providing seeds of Arabidopsis mutants and transgenic lines. We also thank Minna Poranen (University of Helsinki, Finland) for providing Phi6 dsRNA.

## Author Contributions

Conceptualization: MH

Methodology: CH, JM, EB, YY, MR, ARS, LEG, LS, MH

Investigation: CH, ARS, LEG, YY, MR, MH

Visualization: CH, ARS, LEG, YY, LS, MH

Funding acquisition: LS, MH

Project administration: MH

Supervision: LS, MH

Writing - original draft: MH

Writing – review & editing: CH, ARS, JM, LS, MH

## Data and materials availability

All data are available in the main text or the supplementary materials.

## Supplementary Materials

Table S1

Figure S1

## Supplemental Data

**Table S1:** Primers used for RT-qPCR.

**Figure S1:** Development of TMV-GFP infection sites in *N. benthamiana* treated with water, poly(I:C), or flg22. Poly(I:C) and, to some extent, also flg22 treatment delays virus movement. Infection sites in leaves treated with poly(I:C) are smaller, and the rate by which these infection sites increase their size is lower than in infection sites in leaves treated with either water (control) or flg22 (supports **Figure 1A** and **B**).

